# Effects of alertness on perceptual detection and discrimination

**DOI:** 10.1101/2023.03.21.533623

**Authors:** Yanzhi Xu, Martijn Wokke, Valdas Noreika, Corinne Bareham, Sridhar Jagannathan, Stanimira Georgieva, Caterina Trentin, Tristan Bekinschtein

**Author notes:** Corresponding author: Yanzhi Xu.

## Abstract

The level of alertness fluctuates throughout the day, exerting modulatory effects on human cognitive processes at any moment. However, our knowledge of how alertness level interacts with specific cognitive demands and perceptual rules of the task is still limited. Here we use perceptual decision-making paradigms to understand how alertness modulates the detection of a stimulus and the capacity to discriminate one stimulus from another. We analyzed data from four different experiments (113 participants in total): 1 - auditory masking detection; 2 - sensorimotor detection; 3 - auditory spatial discrimination; and 4 - auditory phoneme discrimination, and examined the performance of participants during the natural transition from awake (high alertness) to drowsy (low alertness). First, we fitted psychometric functions to the hit rates across different conditions of difficulty for EEG-defined high and low alertness metastable states, respectively. Second, we performed modelling of slope and threshold for the fitted curves as well as signal detection theory measures of perceptual sensitivity (d’) and response bias (criterion). We found lower detection and discrimination sensitivity to stimuli as alertness level decreases, signalled by a shallower slope of the sigmoidal curve and a lower d’, while the threshold increases slightly and equivalently across experiments during lower alertness. There was no change in the criterion to make the decision during the transition. These results suggest that reduced alertness generally decreases the quality of perceptual decision-making. Zooming in, we observed that the decrease in sensitivity measured by slope was stronger for discrimination than for detection decisions, indicating that lower alertness impairs the precision of decisions in discriminating alternatives more than identifying the presence of a stimulus around the threshold. Taken together, these results suggest that alertness has a common effect on perceptual decision-making and differentially modulates detection and discrimination decisions.

## 1 Introduction

The heterogeneous interactions between the levels of alertness and different cognitive processes are in need of systematic study (Goupil and Bekinschtein, 2012; Mediano et al., 2021). Research on this interplay is important not only to understand human consciousness and cognition but to improve psychological and physiological experiments where the fluctuating levels of alertness become a source of noise that should be measured and controlled. Furthermore, there are health, well-being, and societal implications arising from this interaction such as in shift workers and long-distance drivers, who take decisions based on impoverished behavioural parameters. Among all cognitive processes, perceptual decision-making has been seen as a window to human cognition (Shadlen and Kiani, 2013) and can be easily assessed behaviourally with detection or discrimination tasks. Here, we examine the interaction issue by looking at how the level of alertness affects perceptual decision-making in four independent experiments.

### 1.1 Decision-making processes

Decision-making is a core component of human cognition involving weighing possible choices or behaviours and making judgments based on complex information from the environment and internally present decision rules. People make numerous decisions in daily life, which can be economic, political, ethical, etc. Among these, one simple form is ‘perceptual decision’, the elementary (binary) judgement on sensory perception (O’Connell et al., 2018). Perceptual decision-making is the process of detecting, discriminating, and categorizing information from the senses (Hanks and Summerfield, 2017).

Two commonly used types of experiments in the study of perceptual decision-making are detection and discrimination tasks. In a detection task, a target stimulus is presented with a certain probability and the subjects are required to detect the presence of the target. Subjects are asked to respond ‘yes’ or ‘no’ based on the information they perceive. The hit rate, the percentage of ‘yes’ responses when the stimulus is presented, is usually used to measure performance. In comparison, there are two stimuli in a discrimination task. Subjects are asked to make judgments between two stimuli depending on the signal strength they perceive for each alternative. The accuracy, the percentage of correctly choosing one stimulus when it is presented, is the primary behavioural measure of discrimination.

The hit rate and accuracy are general behavioural measures, which are the manifestation of a series of inner processes, including sensory processing, stimuli recognition, decision-making processing and execution of motor commands. The discrepancy in processing surely exists between detection and discrimination tasks, however, these two types of tasks share the common elements of decision formation that we are specifically interested in this paper. One can consider detection as a comparison between a stimulus and noise while discrimination as comparing between two stimuli. The common decisionmaking stage is the accumulation of sensory evidence until reaching some threshold level (or criterion) and triggering the commitment to a decision (Gold, Shadlen, et al., 2007; Samaha et al., 2020). Empirically, research has shown that contrast thresholds in yes/no detection and orthogonal discrimination tasks are indistinguishable and the two tasks are functionally equivalent (Smith and Ratcliff, 2009; Thomas and Gille, 1979).

To explore perceptual decision formation, models in perceptual psychophysics have been deployed, including psychometric functions and signal detection theory (SDT). A commonly used psychometric function is the sigmoidal function which depicts the relationship between signal intensity and performance (Del Cul et al., 2007; Noreika, Canales-Johnson, et al., 2020; Sandberg et al., 2011). Such relation is further characterized by threshold and slope, two parameters in the sigmoidal function model. Slope reflects the precision of decision, and threshold shows individuals’ perceptual bias (Prins et al., 2016). Another classical model, SDT, better captures the distinction between the perception of sensory evidence and the criterion for making a judgment (Green, Swets, et al., 1966). In the framework of SDT, participants compare criterion with a decision variable which arises from the representation of evidence when making a perceptual decision. The decision is a stage that connects the representation of stimuli to the subject’s behaviour. By using SDT, we can not only deduce the underlying representation of evidence from behaviour (Gold, Shadlen, et al., 2007) but also further examine the decision process itself (Shadlen and Kiani, 2013).

### 1.2 Fluctuation in alertness

As we fall asleep each night, we gradually lose consciousness of the world and ourselves until entering sleep. Low alertness is a state between wakefulness and sleep where people change from being aware and awake in ‘full consciousness’ and capable of responsiveness (Bekinschtein et al., 2009) to reduced responses to stimuli and increased arousal threshold (Goupil and Bekinschtein, 2012). Low alertness happens not only before sleep onset at night but also frequently during the daytime (Carrier and Monk, 2000; Goel et al., 2011). Few studies have concentrated on fluctuations in alertness level and their relationship with cognitive processes (Goupil and Bekinschtein, 2012) despite the pervasiveness of alertness changes throughout the day.

The transition from high to low alertness does not occur in a moment but is a continuum of changes. The complex process of falling asleep involves the convergence of physiological, EEG and behavioural dynamics (Ogilvie, 2001). In addition to physiological changes like reduced heart rate and respiratory activity, one outstanding feature of lower alertness is the change in the frequency spectrum of EEG signals. According to the Hori scoring system that assesses the depth of drowsiness (Hori et al., 1994), the electrical brain activity is dominated by the alpha band during relaxed wakefulness. As people get into mild drowsiness, the alpha wave trains become less frequent and the EEG activity may even become flat. Instead, another oscillation with a higher amplitude, theta wave, emerges. When people enter deeper stages in the transition, vertex sharp waves, spindles and K-complexes appear successively.

At the behavioural level, a more obvious characteristic of sleep onset is that subjects gradually lose the ability to respond to external stimuli, including the diminished performance of accuracy, reaction time and response rate (Bareham et al., 2014; Noreika, Canales-Johnson, et al., 2020; Ogilvie, 2001). Researchers have used short reaction time to indicate wakefulness and longer reaction time to mark decreasing alertness (Ogilvie et al., 1991). As people go deeper into the metastable states in the transition their responsiveness drops or ceases (Harsh et al., 1994), and unresponsiveness is taken as evidence of later stages of the N1 and entering true N2 sleep (Ogilvie, 2001). These behavioural dynamics are also correlated to EEG changes during the transition to sleep. Reaction time increased along Hori stages, while response cessation is usually found during later Hori stages (Hori et al., 1994; Liberson and Liberson, 1966).

### 1.3 Current study

Decision-making performance is affected by the strength of signal intensity, with higher intensity leading to a higher probability of entering participants’ awareness in detection tasks or a higher probability of discerning the identity of the stimuli in discrimination tasks (Koch and Preuschoff, 2007; Marcel, 1983; Sergent and Dehaene, 2004). This reflects how externally driven noise impairs our cognitive performance. On the other hand, inner noise such as a lower alertness level also exerts influences on cognitive processes. Previous research introduced both internal and external noises and revealed how a person’s alertness level *per se* influences perceptual decision-making with varied signal intensities (Noreika, Canales-Johnson, et al., 2020; Noreika et al., 2017). However, it is still unknown if the alertness fluctuation exerts a common effect on detection and discrimination decision-making or if it interacts with the specific cognitive demands and perceptual rules of the task at hand.

In this study, we are interested in the effect of alertness level on decision-making performance in simple detection and discrimination tasks. In other words, how the detection and discrimination performance changes as people go from fully awake to low alertness metastable states. To this end, we analyse data from four experiments, including two detection tasks (auditory masking and sensorimotor experiments) and two discrimination tasks (spatial attention and phoneme discrimination experiments), and use both psychometric function and SDT to investigate the change in the properties of decision-making in detection and discrimination tasks during the natural transition from wakefulness to low alertness.

The methods we apply in this study are as follows. First, all trials in each experiment are divided into high alertness or low alertness based on 4 different methods: a) an automatic micro-measures algorithm (Jagannathan et al., 2018) that uses EEG data of 4 seconds before the stimulus onset to capture the variance and coherence features of the EEG frequency space and recognize the patterns of elements like vertex, K complex, and spindles; b) *θ/α* ratio on the 2-second pretrial signal; c) 45 percent median split on fast and slow reaction times (RTs); d) proportion of missed responses within 10 consecutive trials. Second, changes in performance indices between alertness levels are investigated. At each alertness level, the hit rate of target stimuli with varied intensities is modelled per participant using a classic psychometric sigmoidal function. Then slope and threshold obtained from the sigmoidal function for each participant are compared between high and low alertness within each experiment. Further, we apply SDT to analytically decouple perceptual sensitivity from response biases, to characterize the mechanisms underlying the behavioural changes. Last but not least, we use multilevel modelling (MLM) to test whether different perceptual demands have an impact on the relationship between alertness and decision-making performance.

The hypotheses tested in this study stem from the preregistration (https://osf.io/3p4rf). We hypothesize that alertness level has a common effect on detection/discrimination for all four experimental settings: as people enter lower alertness, their precision of decision (slope of the sigmoidal curve) decreases, while their threshold (the stimulus intensity at the inflexion point of the sigmoidal curve) may increase. Relative hypotheses from the preregistration are as follows:

H1: We hypothesize that the proportion of correct responses to detect or discriminate between stimuli increases with stimulus intensity following a sigmoidal curve, regardless of conscious state.
H2: As people become drowsy, their general sensitivity (the slope of the sigmoidal curve) decreases (as seen in Noreika, Kamke, Canales-Johnson, Chennu, Bekinschtein, et al., 2020), which may be attributed to reduced perceptual sensitivity (d’). Specifically:

a. as alertness decreases, detection sensitivity decreases.
b. as alertness decreases, discrimination between stimuli sensitivity decreases.
H3: As people become drowsy, their threshold (the stimulus intensity at the inflection point) may increase. This is determined by an increase in the criterion relative to the sensory distribution.

a. drowsy states will increase the detection threshold.
b. drowsy states will increase the discrimination threshold.
H4: Common effect of alertness: Alertness level has a main effect on detection/discrimination for all four experimental settings, both in threshold (increase) and slope (decrease).

## 2 Methods

### 2.1 Participants and experiments

Four studies are analyzed to explore the interaction between the level of alertness and perceptual decision-making performance (Fig. 1). The first two studies are detection tasks and the other two are discrimination tasks. These are explained in detail as follows but in short, they are auditory detection of a target masked by white noise, kinaesthetic detection of a muscle movement induced by transcranial magnetic stimulation (TMS), auditory spatial discrimination, and phoneme morphing discrimination. For all experiments, easy-sleeper participants were seated comfortably with their eyes closed in a dimmed room and were encouraged to relax during these tasks, allowing the low alertness state to emerge while responding to the stimuli or cues. For the two discrimination tasks, in addition to the above drowsy session, there was an awake session when participants were seated upright with lights on and instructed to stay awake. High and low alertness were classified within the drowsy session for detection tasks and were determined between awake and drowsy sessions in discrimination tasks. For both high and low alertness levels, we assess participants’ performance according to varied stimuli intensities. The difference in performance among experiments is further examined in order to test a common effect of alertness level on cognitive processing.

**Figure 1:**
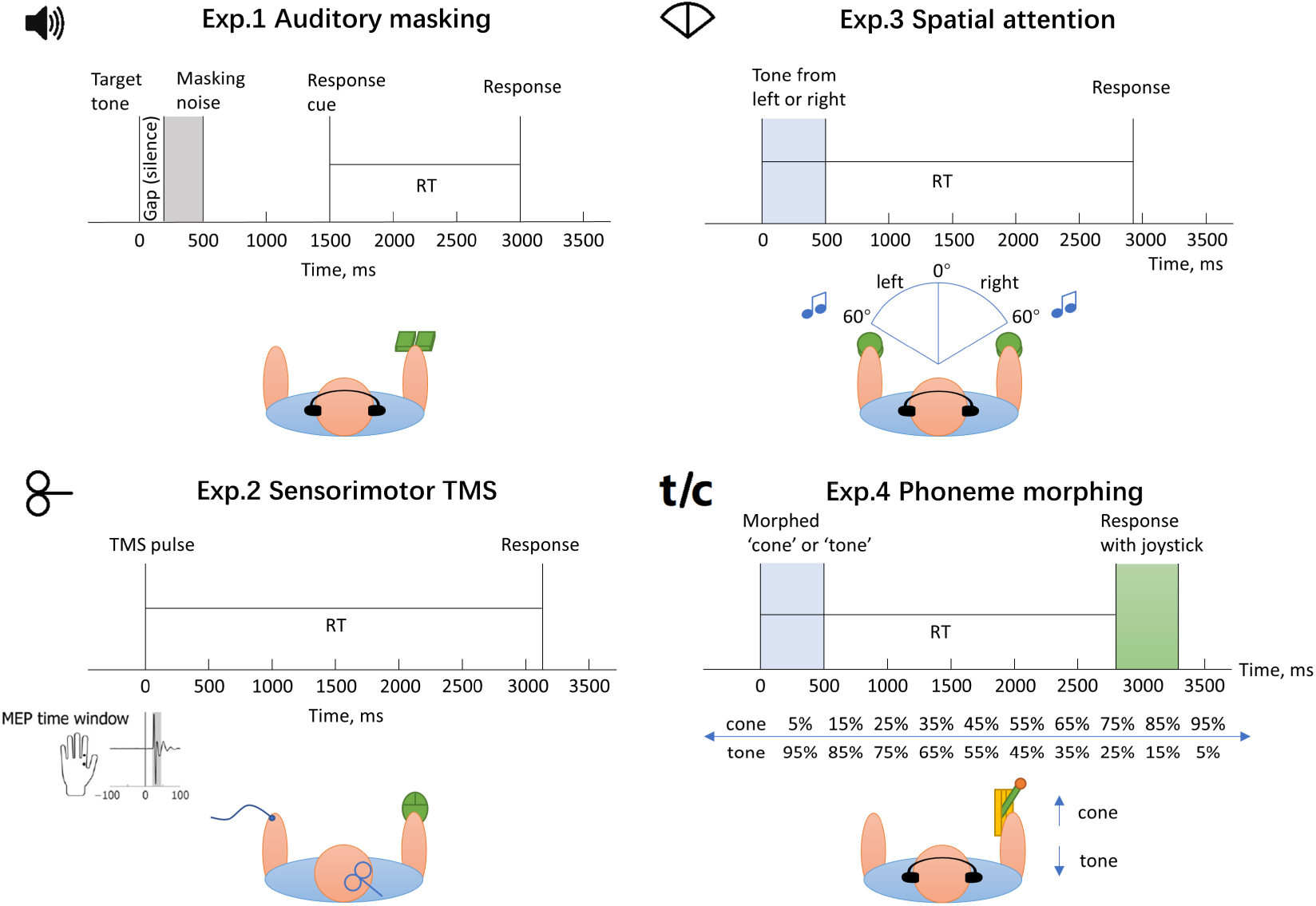
Experimental design. The experimental design of four tasks is shown (the left two are detection tasks and the right two are discrimination tasks). The first used a target tone masked by a noise with a varying gap. Participants were asked to respond to whether they heard the target tone after a response cue. The second experiment was a sensorimotor task, different from the other three tasks using auditory modality. TMS with different intensities around individual thresholds was applied to the right motor cortex of participants. They responded using their right hand to indicate whether they felt a twitch in the left hand where motor-evoked potentials (MEPs) were recorded as well. In the third task, participants were presented with the target tones that fell on their left or right side and responded to indicate the location of the target with left or right buttons. The last task used the morphed phoneme of ‘cone’ and ‘tone’ as the target and participants were asked to move a joystick to indicate the word they heard. The distance they moved the joystick represented their confidence in the answer. By including these tasks, we covered a wide range of perceptual decision-making processes in the testing of the effects of alertness levels.

Experiment 1 (auditory masking detection https://www.biorxiv.org/content/10.1101/155705v3): This data was collected to investigate the influence of the wakefulness state on auditory detection, and the basic threshold and slope but not SDT results were already published (Noreika, Canales-Johnson, et al., 2020). 56 adults were recruited to participate in an auditory detection task through the electronic volunteer database of the MRC Cognition and Brain Sciences Unit at the University of Cambridge. They were screened using the Epworth Sleepiness Scale (Johns, 1991) with a minimum sleepiness score of 7 in order to be likely to become drowsy in the experiment. After the screening, 31 participants remained in the study (9 male; mean age 27.4; age range 20-39). During the 2-hour experiment, participants relaxed and were presented with a target sound (10 ms; 1000 Hz; 2.5 ms fade-in and fade-out; attenuation: −24 Db) and a following masking sound of white noise (300 ms; 707.1-1414.2 Hz; 5 ms fade-in and fade-out; attenuation: 0 Db), between which the duration was varied near the individual threshold (Fig.1: Exp.1). The individual threshold (the duration between target sound and masking noise) was estimated before the experiment. Participants were asked to report if they heard a target sound by hitting one of two buttons after a response cue. No response within 6 seconds from the onset of the target stimulus was defined as unresponsive. There were 11 conditions of the interval between the target stimulus and the noise (0, 0.25, 0.5, 0.75, 0.875, 1, 1.125, 1.25, 1.5, 1.75, 2 * individual threshold). Additional catch trials were also performed where there was no target sound before the mask. On average, 501 trials were carried out per participant (SD=65, Min=361, Max=604). EEG signals were recorded during the whole experiment. The experimental protocol was approved by the Cambridge Psychology Research Ethics Committee, and participants received a remuneration of £30 for the study.

Experiment 2 (sensorimotor detection to TMS https://www.sciencedirect.com/science/article/pii/S1053811920307916): The experiment was conducted to explore how cortical evoked responses to TMS are modulated by alertness level, and the basic slope and threshold but not SDT results were published (Noreika et al., 2017; Noreika, Kamke, Canales-Johnson, Chennu, Bekinschtein, et al., 2020). 20 right-handed participants (7 male; mean age 23.7; age range 21-33) attended this experiment. They were also screened with the Epworth Sleepiness Scale (Johns, 1991). The mean ESS score was 9.4 (SD=4.3). 17 participants remained in the analysis after excluding those with too low responding rates. TMS was applied to the right motor cortex with the output intensity varying around the individual resting motor threshold (−20%, −15%, −10%, −5%, 0%, +5%, +10%, +15%, +20%), which was determined using a criterion of ≥ 50*μ*V motor evoked potential amplitude in at least five out of ten consecutive trials (Ikoma et al., 1996; Rossini et al., 1994; Samii et al., 1996). Meanwhile, Surface Electromyography (EMG) from the first dorsal interosseous (FDI) of both hands and scalp EEG activities were recorded during the experiment (Fig.1: Exp.2). Participants were instructed to respond whether they felt a twitch or a touch in their left hand after each TMS pulse by clicking mouse buttons with their right hand. No response within 6 seconds after TMS was considered an omission. 520 trials were performed for each participant with inter-pulse intervals of 8.5-10.5 seconds. The experimental protocol was approved by the Medical Research Ethics Committee of The University of Queensland and each participant received a remuneration of $30.

Experiment 3 (spatial attention discrimination https://www.jneurosci.org/content/42/3/454): In this experiment, complex harmonic tones were used to investigate the effect of alertness level on spatial attention. Psychometric and SDT results of the experiment were published (Jagannathan et al., 2022). 41 right-handers were recruited for this study. After discarding invalid data due to technical problems or not following instructions, 32 participants (14 male; 24.46 ± 3.72 years old) remained in the analysis, among which 29 of them had a sleepiness score ≥ 7 (easy sleepers) and 3 had a sleepiness score ≥ 4 of the Epworth Sleepiness scale (Johns, 1991). All participants attended two sessions (awake and drowsy sessions). The awake session lasted about 8 minutes and was comprised of 124 trials. The drowsy session lasted 1.5-2 hours and included 740 trials. Both sessions contained the same auditory tone localization task in which participants were presented with complex harmonic tones from left 59.31° to right 59.31° of their spatial midline (Fig.1: Exp.3). 12 steps of 4.01° were used between 11.17° and 59.31° in both left and right directions, and smaller steps of 1.86° were used between left 11.17° and right 11.17°. Participants were instructed to report the direction of the tone (left/right) with a button press. The response window was 5 seconds. EEG was also recorded during the whole experiment. Participants received £30 for taking part in the study.

Experiment 4 (phoneme morphing discrimination): 33 right-handed participants completed the EEG experiment and entered a lower alertness state with enough trials for analysis. They were presented with a mixture of morphed spoken words ‘tone’ and ‘cone’. Ten different percentages of morphing between the two words were used as conditions of difficulty (Fig.1: Exp.4). The participants reported whether they had heard the word ‘tone’ or ‘cone’ and their confidence in the answer by moving a joystick. The direction of the movement (forwards or backwards) indicated the choice of the word and the extent to which they moved indicated confidence. No feedback was provided throughout the task. The experiment consisted of a 12-minute awake session (160 trials) and a roughly 85-minute drowsy session (480 trials). During the awake session, the response window is 2.5 seconds and inter-stimulus intervals varied between 2.5 and 4 seconds. In the drowsy session, the response window is 4 seconds and inter-stimulus intervals ranged between 4 and 8 seconds. Participants were woken up when they had missed 3 consecutive responses. The study was approved by the ethical committee of the Medical Research Council for the Cambridge Brain Sciences Unit.

### 2.2 EEG acquisition and preprocessing

Experiment 1 (auditory masking): 128-channel EEG data were recorded at a sampling rate of 500 Hz using the Net Amps 300 amplifier (Electrical Geodesics Inc., Oregon, USA). After excluding channels over forehead, cheeks and neck, 92 channels were retained in the analysis. Raw data have been filtered between 0.5 Hz and 40 Hz, re-referenced to the average, and epoched between −4000 ms and 6000 ms relative to the onset of the target sound.

Experiment 2 (sensorimotor TMS): For this TMS-EEG combined experiment, 64-channel EEG data were recorded using the BrainAmp MR Plus amplifier, TMS BrainCap, and Brain Vision Recorder (v1) software (Brain Products; Gilching, Germany). The recording was down-sampled to 500 Hz and epoched between −4000 ms and −10 ms relative to each TMS pulse for calculating EEG spectral power before TMS.

Experiment 3 (spatial attention): During both awake and drowsy sessions, 128-channel EEG data were recorded at a sampling rate of 500 Hz (Electrical Geodesics Inc., Oregon, USA). Channels over forehead, cheeks and neck were excluded, retaining 92 channels in the analysis. Raw data have been filtered between 1 Hz and 40 Hz and epoched between −200 ms and 800 ms relative to the onset of the tone stimuli as post-trial epochs. Additionally, data were epoched between −4000 ms and 0 ms to the onset of the stimuli in the drowsy session for classifying the alertness level.

Experiment 4 (phoneme morphing): EEG data were recorded at a sampling rate of 1000 Hz by using 128-channel HydroCel Sensors and a GES300 amplifier (Electrical Geodesics Inc., Oregon, USA), and 92 channels were retained in the analysis. Raw data have been filtered between 0.5 Hz and 40 Hz, re-referenced to the average of all electrodes, and epoched between −200 ms and 2002 ms relative to the onset of the target sound. Drowsy session data were also epoched between −4000 ms and 0 ms to the onset of the stimuli for classification of alertness levels.

Furthermore, for all experiments, independent component analysis (ICA) was performed to remove artefacts and noisy channels were interpolated.

### 2.3 Measures of alertness

The first step in the analysis is to classify the alertness level of subjects during the experiments. There is no single widely accepted measure of lower alertness in cognitive experiments. Therefore, we used a combination of four methods to validate and assess the degree of methods employed to define lower alertness: 1) micro-measures based automatic classification of alertness levels based on multiple parameters including EEG power/coherence and spatio-temporal signatures (Jagannathan et al., 2018); 2) relative change in *θ/α* spectral power ratio; 3) relative change in reaction time length; 4) proportion of missed trials. In detection tasks, trials were divided into ‘high alertness’ and ‘low alertness’ using the above methods. In discrimination experiments, since they contained extra awake sessions, we classified all trials in awake sessions as ‘high alertness’ and lower alertness trials in drowsy sessions as ‘low alertness’.

#### 2.3.1 Micro-measures

Micro-measures algorithm is an automated method to detect micro variations in levels of alertness in EEG experiments under eye-closed settings (Jagannathan et al., 2018). It applies a support vector machine (SVM) and individual element detectors on the 4-second pretrial EEG data to classify each trial into different alertness levels. Specifically, two major steps were conducted to generate the micro-measures algorithm. First, predictor variance and coherence features were computed and then used to classify data into ‘awake’ and ‘drowsy’ using the SVM. Individual alpha band range (individual alpha peak ±2 Hz) was used in the computation. Second, the drowsy data were further examined using individual element detectors to detect vertex, K-complex, and spindles. Then another SVM was used to detect true spindles based on the variance and coherence features. Data with vertex, K-complex, and true spindles were identified as ‘late drowsy’ while other drowsy data were marked as ‘early drowsy’. In our analysis, both ‘late drowsy’ and ‘early drowsy’ trials were categorized into low alertness and further compared with high alertness state (awake).

#### 2.3.2 EEG *θ/α* power

Previous research has shown that lower alertness can be characterized by increased theta band power and decreased alpha band power (Hori et al., 1994), or an increased ratio of *θ/α* (Bareham et al., 2014; Noreika, Canales-Johnson, et al., 2020). Here, we computed a ratio of *θ/α* power based on the pretrial data, according to which we split the data into equal proportions of high alertness and lower alertness trials for each participant. More specifically, data from 2-second pretrial epochs with respect to the onset of stimuli were used to compute the spectral power of EEG frequency oscillations. *θ* and *α* power were extracted respectively from the results of complex Morlet wavelet convolution (M. X. Cohen, 2014), in which nine frequencies logarithmically increased between 4 Hz and 12 Hz and the number of wavelet cycles was 10. Convolution results were first averaged over −2 s to 0 s. Then data of 4-7 Hz were averaged to obtain *θ* power and data of 8-12 Hz were averaged to calculate *α* power for each electrode and each trial. The *θ/α* ratio was then averaged across all electrodes. In this way, we obtained a single *θ/α* value for each trial. Data for each participant were divided into equal proportions of lower 45% and higher 45% *θ/α* power ratio as high alertness and low alertness, excluding 10% intermediate trials.

#### 2.3.3 Reaction times (RTs)

The alertness level can also be measured by RT and RT variability (Bareham et al., 2014; Ogilvie, 2001). Studies show that people respond slower as they become drowsy compared to awake (Hori et al., 1994; Ogilvie et al., 1989). To apply the RTs measure, the same criterion of equal proportions was used as *θ/α* power ratio. Trials with 45% shortest RTs were labelled as high alertness and trials with 45% longest RTs were marked as low alertness for each participant. The advantage of the RTs measure is that it reveals the alertness level at the exact time of response rather than seconds before the response as the micro-measures and *θ/α* ratio measure. However, some research shows no association between RT and lower alertness (Baulk et al., 2001). So we cautiously use the RTs measure as a complementary tool.

#### 2.3.4 The proportion of omissions

As alertness decreases, responses of subjects become progressively intermittent (Lagarde and Batejat, 1994; Makeig et al., 2000; Ogilvie et al., 1989). The lack of response to the stimuli could also be used as a measure of sleepiness (Williams et al., 1959). Previous research on lower alertness inspected the sequence of omissions and marked omission as a lack of response within a 6-s window after stimulus presentation (Comsa et al., 2019). In this paper, we inspected the proportion of missed responses to indicate subjects’ alertness level. Behavioural data were divided into blocks of 10 trials and the number of unresponsiveness within each block was counted. Blocks with 2 or more omissions were marked as low alertness and blocks with no omissions were marked as high alertness (Bareham et al., 2014). Blocks with 1 omission were intermediate trials and were excluded from further analysis. This method of alertness measurement is therefore even more diluted in time.

### 2.4 Signal detection theory analysis

Signal detection theory (SDT) has been widely used in the analysis of detection and discrimination experiments (Macmillan and Creelman, 2004). The significant contribution of SDT is that it disentangles the encoding process and decision-making process by computing two parametric statistics: the perceptual sensitivity (d’) and criterion (C) (Green, Swets, et al., 1966). The sensitivity d’ describes the sensory perception of the stimuli while C reflects the boundary participants set. When sensory evidence crosses that boundary, a response is given that a stimulus is present (or that an alternative stimulus is present). Criterion could capture the decision bias when people make a response. Here, perceptual sensitivity and criterion were calculated based on behavioural performance (Equation 1 and 2) for each participant and each level of alertness and further modelled by alertness and experiment later.

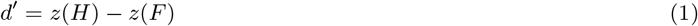

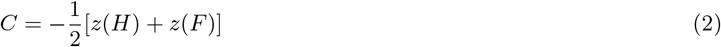

where *H* represents the hit rate, *F* represents the false alarm rate and *z* stands for transformation to a *z* score.

More specifically, for the auditory masking task, the false alarm rate was calculated based on catch trials and the hit rate was obtained from the pooled performance across different conditions of difficulty. For the TMS task, the weakest TMS pulse was used to calculate the false alarm rate since there were no catch trials in this experiment. To make the parameters comparable among detection and discrimination tasks, we treated one stimulus in the discrimination tasks as the target and another stimulus as noise. Therefore, the weakest condition for the target stimulus was used to derive the false alarm and all other conditions were pooled together to get the hit rate in the two discrimination experiments. For the spatial attention task, the ‘left’ responses were considered as hits, because participants made more errors for left-sided stimuli than for right-sided stimuli as their alertness level decreased (Jagannathan et al., 2022) and we would like to avoid this left bias when calculating the false alarm rate in SDT analysis. For the phoneme morphing discrimination task, we randomly used ‘cone’ as the target stimulus and used 5% ‘cone’ condition to obtain the false alarm rate^1^.

### 2.5 Psychometric curve fitting

To describe the change in detection and discrimination with different alertness levels, we fitted the psychometric curves to the hit rates of target stimuli with different intensities in high and low alertness trials, respectively. Since the birth of SDT (Green, Swets, et al., 1966) and its use in experimental psychology, several methods have been developed to fit the performance curves. One way is to model psychometric functions with SDT (Prins et al., 2016). Since d’ and the proportion of correct responses could be mutually converted, we could construct a function that describes the relationship between stimulus intensity and d’, and then further convert it to behavioural performance. Another way of fitting a psychometric curve is to use sigmoidal functions that describe the relationship between stimuli and performance directly, with parameters of slope and threshold (Prins et al., 2016). Sigmoidal functions commonly used include cumulative normal distribution, logistic and Weibull functions. The Weibull function is given as:

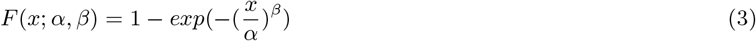

where *α* is threshold corresponding to *F*(*x* = *α*; *α, β*) = 1 – *exp*(– 1^*β*^) ≈ 0.632 and *β* is slope. We used the Weibull function because the former two are inappropriate when stimulus intensity x=0 represents an absence of signal (Prins et al., 2016). The psychometric function is further formulated as:

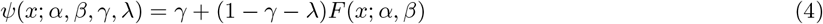

where *γ* corresponds to the guess rate (lower asymptote), and λ corresponds to the lapse rate (upper asymptote).

We fitted the performance curves using the SDT method and Weibull function with the Palamedes toolbox for Matlab (Prins and Kingdom, 2018), and the latter had a better fitting with our data so we show the outcomes from Weibull fitting in this paper. One possible reason is that there are not enough trials to produce a reliable estimation of d’ in each condition of difficulty for SDT psychometric fitting. Besides the choice of the fitting function, the setting of guess rate γ and lapse rate λ may also influence the fitting outcomes (Green, Swets, et al., 1966; Wichmann and Hill, 2001). Guess rate is the chance that participants give a positive response when they do not detect the stimuli, and the lapse rate corresponds to the percentage of errors made for most detectible stimuli (Gold and Ding, 2013). We first combined performance in high and low alertness and fit the curve to find a common guess rate and lapse rate for different alertness levels, and then fixed the guess rate and lapse rate as the common ones in the separate fittings of performance in high and low alertness (except for the lapse rate in spatial attention task due to the left bias: we set the lapse rate to be different for high and low alertness to better fit the performance). In this way, different alertness levels shared the same guess rate and lapse rate settings so that we could focus on the comparison between slope and threshold. If we see effects when using the same guess rate and lapse rate for different alertness levels as we have found in this paper, we could expect larger effects when we free the restriction of parameters. In addition, we fitted the individual psychometric curves using different settings, which were individual best parameters, group best parameters and fixed parameters of 0.05 for guess rate and lapse rate. In the main text, we show fitting results using individual best guess and lapse rates. Results using group best rates (group fitting) and 0.05 (strict fitting) are shown in the supplementary information. The different asymptote values did not influence the pattern of results and we could be more confident in the findings of alertness’s effects on slope and threshold.

### 2.6 Multilevel modelling

To investigate the common effect of alertness on detection and discrimination decision-making, MLM was used on the above behavioural measures (slope, threshold, d’ and criterion). The difference between high and low alertness was first illustrated using raincloud plots (Allen et al., 2021) and statistically compared within each experiment. Next, MLM was performed on the aggregated data of four experiments using the lmer function from the lme4 package in R, to compare the difference in the behavioural performance between high and low alertness across different experimental settings. Subject identity was used as a random factor, and alertness level and experiment were used as fixed factors. Models with different combinations of fixed factors (alertness alone, experiment alone and both alertness and experiment) were compared using Akaike Information Criterion (AIC), Bayesian Information Criterion (BIC) and the negative log-likelihood. The F test for the complete model (alertness * experiment) was further conducted by using the anova function.

## 3 Results

In the following sections, We present statistical and modelling results on four behaviour measures: slope and threshold from psychometric curve fitting, and d’ and criterion from SDT. We concentrate on the effects of alertness captured by the micro-measures method and compare them, in the text, to the other three alertness split methods (see supplementary information for further results and specific figures). The complementary analyses on slope and threshold using different curve fitting parameters and analyses on slope using integrated sessions in discrimination tasks are also discussed, and detailed results are shown in the supplementary information.

### 3.1 The precision of decision decreases in low alertness states

For both states of alertness, performance follows a sigmoidal curve. Critically, however, with decreased alertness, the curves are markedly shallower (lower slope) as compared to the high alertness across all experiments (see Fig. 2). This suggests that participants are less sensitive to the change in stimulus intensity as they become drowsy. In other words, their precision of decision is reduced in lower alertness. To formally test this (hypothesis H2 in the preregistration), we compared the distributions of slopes between different alertness states for each experiment and performed MLM on the aggregated data of all tasks. The main founding is that alertness level has a strong effect on slope (*F*_(1,80)_ = 51.228, *p* < 0.001) as revealed by the analysis of variance in MLM (Table 2).

**Figure 2:**
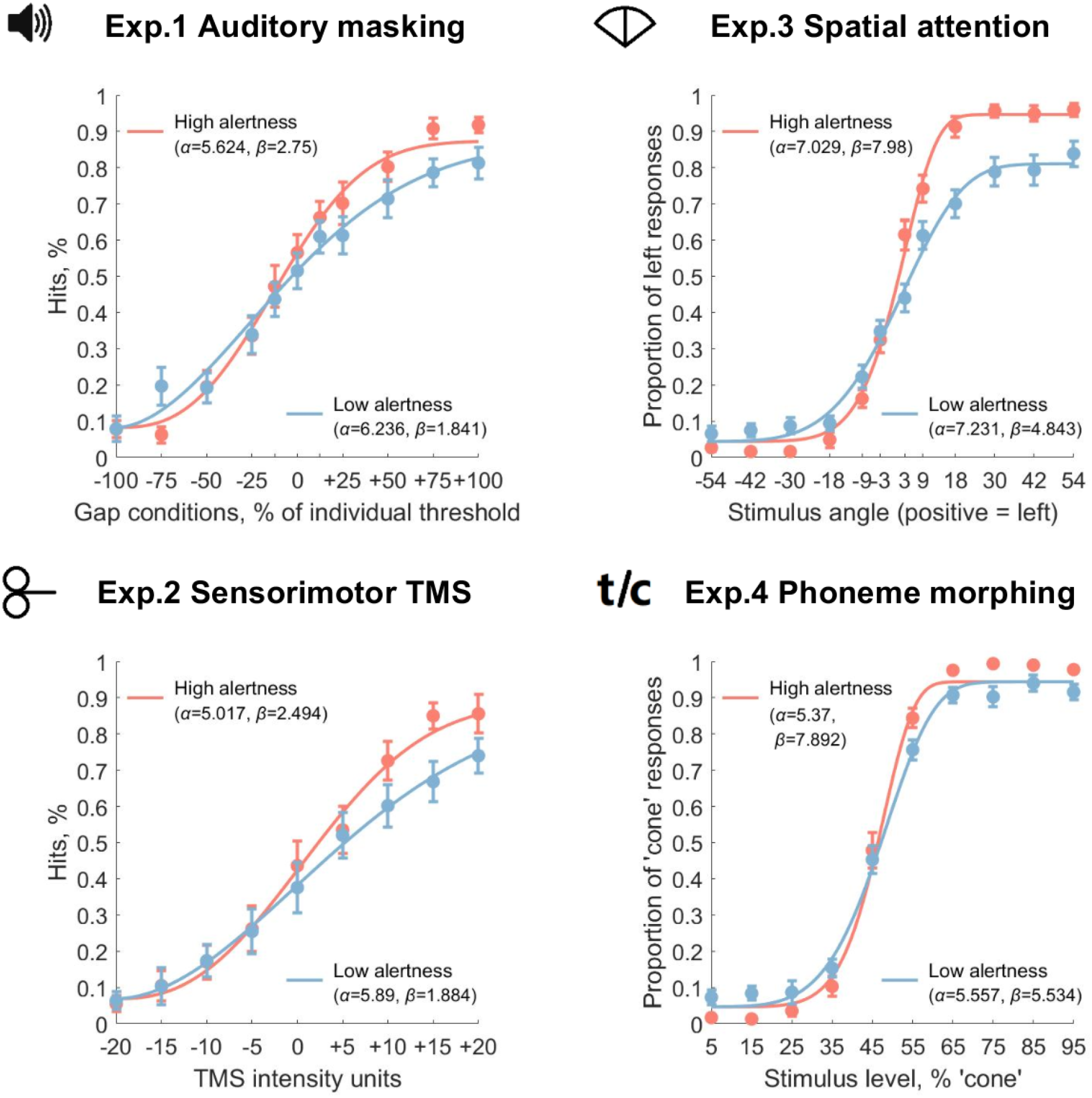
Psychometric curve fitting. High and low alertness sigmoidal curves are shown for all decision-making tasks, fitted with the Weibull function for each alertness level. The error bars indicate variations across individual subjects. Alertness levels are defined by micro-measures method. Slope (β) shows a systematic decrease with low alertness and the effect interacts with experiment settings (see Table 1 and Table 2 for MLM results). Threshold (α) shows a reliable increase with lower alertness (Table 3 and Table 4). See text for more details.

In detail, in the comparison of slope in each task, we found a medium to large effect of reduced slope for lower alertness level (Fig. 3). The experiment on spatial attention showed a larger effect than the other three experiments. Furthermore, slope shows a larger variance in the high alertness level (significant for Exp. 1 (*F*_(16,16)_ = 3.193, *p* = 0.026) and Exp. 4 (*F*_(22,22)_ = 3.783, *p* = 0.003), and marginally significant for Exp. 2 (*F*_(15,15)_ = 2.546, *p* = 0.08) and Exp. 3 (*F*(_23,23)_ = 2.075, *p* = 0.087)), pointing to a larger individual difference in sensitivity when people are awake as compared to low alertness. The long tail in the high alertness state indicates that some participants have much better performance than the majority.

**Figure 3:**
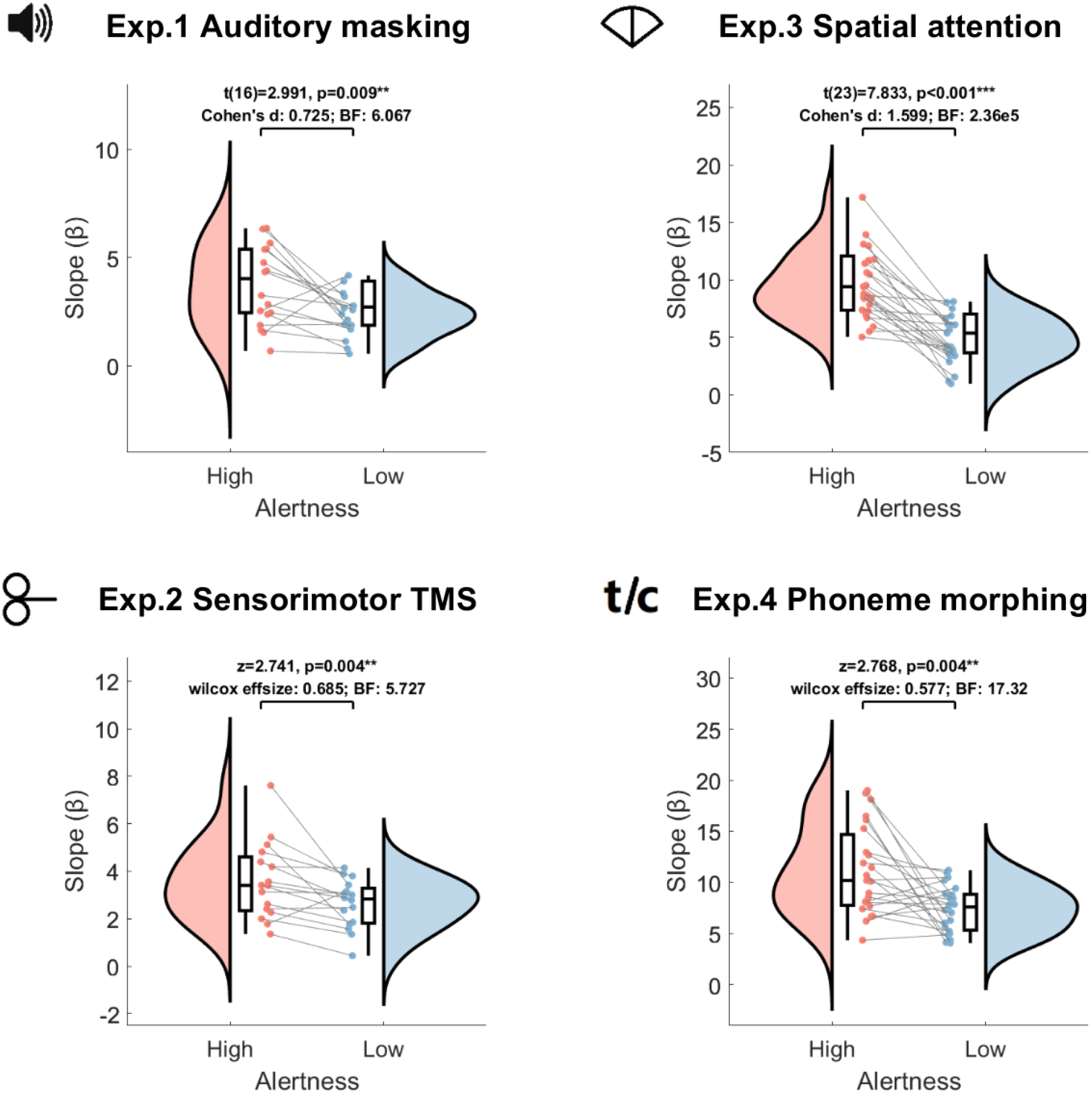
Distributions of slope. Distributions of slope are shown for each task and each alertness level (defined by the micro-measures method). Results of the t-test or Wilcoxon signed-rank test, effect size and the Bayes factor are shown in the figure. Slope decreases significantly with alertness level for all tasks. MLM reveals a reliable effect of alertness and an interaction effect between alertness and experiment settings (Table 1 - 2). *Post hoc* analysis (Table S3) shows that the decrease in slope with lower alertness is more pronounced in discrimination than in detection tasks.

We further combined all tasks and performed MLM analysis using participant as a random factor and various combinations of alertness level and experiment as fixed factors to examine the effect of alertness level on slope among different task settings. The winning model (with the smallest values of AIC and BIC and the largest value of negative log-likelihood) comprised both alertness level and experiment factors (Table 1). As pointed out in the introduction of this section, the main effect of alertness level on slope is significant (Table 2). This means that regardless of the different experimental settings, the sensitivity to stimulus intensity in perceptual decision-making is modulated by the alertness level of participants during the task. The factor ‘experiment’ alone also shows a reliable effect on slope (*F*_(3,80)_ = 60.006, *p* < 0.001). As depicted in Fig. 2 and Fig. 3, the curves of discrimination tasks are steeper (larger slope) than that of the detection tasks. The experiment of spatial attention exhibited an asymmetry for left and right stimuli which is consistent with previous findings (Jagannathan et al., 2022).

**Table 1:** Model comparison for slope.

| Model | Parameter | AIC | BIC | Log-likelihood | $p > (x^2)$ |
| --- | --- | --- | --- | --- | --- |
| Null | Fixed: mean, Random: participant ID | 894.39 | 903.62 | -444.20 | - |
| Alertness | Fixed: alertness,<br>Random: participant ID | 856.05 | 868.36 | -424.03 | 2.14e-10 *** |
| Experiment | Fixed: experiment,<br>Random: participant ID | 812.75 | 831.20 | -400.37 | < 2.2e-16 *** |
| Alertness * Experiment | Fixed: alertness * experiment,<br>Random: participant ID | 757.23 | 787.98 | -368.61 | < 2.2e-16 *** |

\*\*\* $p < 0.001$ .**Table 2: Type III analysis of variance table for the alertness \* experiment model on slope**

**Table 2:** Type III analysis of variance table for the alertness * experiment model on slope.

| Model elements | Sum Sq | Mean Sq | NumDF | DenDF | $F$ | $P(> F)$ |
| --- | --- | --- | --- | --- | --- | --- |
| Alertness | 281.77 | 281.77 | 1 | 80 | 51.228 | 3.53e-10 *** |
| Experiment | 90.15 | 330.05 | 3 | 80 | 60.006 | < 2.2e-16 *** |
| Alertness:experiment | 100.99 | 33.66 | 3 | 80 | 6.12 | 8.42e-4 *** |

\*\*\* $p < 0.001$ .

We also observed a reliable interaction effect between alertness level and experiment (*F*_(3,80)_ = 6.12, *p* < 0.001). *Post hoc* analysis (Table S3) shows that the alertness level has a significant effect on slope in discrimination tasks (*p* < 0.001 for both spatial attention and phoneme morphing discrimination task) but not in detection tasks (*p* = 0.737 for auditory masking task and *p* = 0.94 for TMS-induced sensorimotor task). These results not only reveal a general effect of alertness on the sensitivity of perceptual decision-making, but also show that the effect is stronger for discrimination than for detection.

To further test the results obtained from the micro-measures alertness split and look for convergent evidence, we also performed the analysis using the other three measures to categorize high and low alertness levels: *θ/α* ratio, fast and slow RTs, or the proportion of missed responses within 10 trials (Fig. S2). Alertness * experiment is the winning model in MLM analysis for all three ancillary methods as it was for the micro-measures method (Table S1). The results confirm the findings with the micro-measures split method that the slope of the psychometric curve decreases as the alertness level gets lower, and the effect is reliable regardless of experiment type (Table S2). We also find reliable effects of experiment and interaction on slope by using the other three split methods. *Post hoc* analysis reveals that the larger effects of decreasing alertness for discrimination tasks are consistent among different split methods (Table S3). Taken together, the main effects of low alertness and its interaction effects with experiment type on slope are convergent and robust across the methodologies used to define alertness.

Two complementary analyses were performed to confirm these results. First, we used the integrated session (high alert trials during the drowsy session) in the discrimination tasks to compare with the analysis using the awake session. The extra analysis showed similar results for the winning model and the effect of alertness despite more lax conditions of high alertness levels (Supplementary Section 3). The interaction effects are, unsurprisingly, less consistent between methods. Second, to test the independence of our main effects from specific parameter specifications (guess rate *γ* and lapse rate λ) used to construct individual curves, we fitted the curve using a pair of group best *γ* and λ rate and a more strict pair (*γ* = λ = 0.05), respectively, showing that the individual variances in the curve fitting do not impair our main conclusion that alertness level has a common effect on slope across different experiments (Supplementary Section 4).

### 3.2 Detection and discrimination thresholds increase with decreasing alertness

We further characterize the effect of alertness on decision-making by testing its effects on threshold of the psychometric curve, revealing the limit of detection and discrimination ability and allowing us to see whether there is a bias in the participants’ decision-making abilities with the change of alertness level. The comparison between the distributions of threshold in different states shows a small increase in threshold for low alertness (Fig. 4). Paired-sample t-tests show that the increase in threshold is substantial only for the TMS-induced kinaesthetic detection experiment (*t* = 2.51, *p* = 0.027).

**Figure 4:**
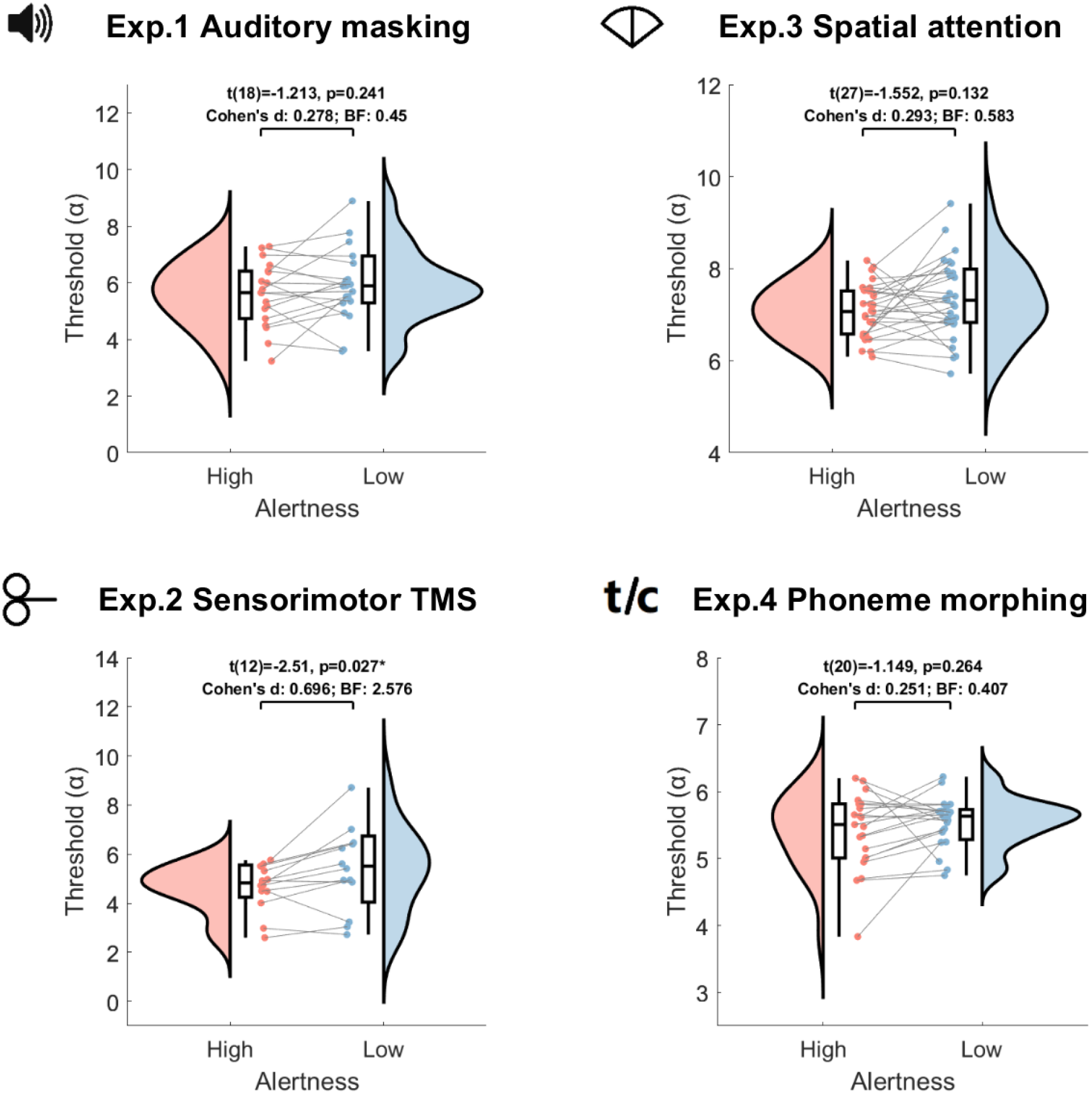
Distributions of threshold. Distributions of threshold are shown for each task and each alertness level (defined by the micro-measures method). Results of the t-test, effect size and the Bayes factor are shown in the figure. Threshold shows a reliable increase in the TMS task and no obvious change in other tasks. MLM including all tasks reveals a reliable effect of alertness level on the threshold (see text and Table 3 - 4).

To test the general effect of alertness on threshold, we also performed the MLM analysis as we did for slope. The alertness * experiment model has the smallest values of AIC and the largest value of negative log-likelihood, and the experiment model has the smallest values of BIC (Table 3). Further comparison shows the alertness * experiment model is the winning model again (*Chi – Squared* = 13.621, *p* = 0.009). The analysis of variance (Table 4) shows that the main effect of alertness is reliable (*F*_(1,81)_ = 12.859, *p* < 0.001), suggesting that threshold of the curves increases as the alertness level goes down as a common pattern of alertness. The factor ‘experiment’ also shows a main effect on threshold (*F*_(3,81)_ = 30.692, *p* < 0.001) while the interaction effect between alertness and experiment is not reliable (*F*_(3,81)_ = 1.435, *p* = 0.239). To check the possibility that the overall effect of alertness is driven by TMS task alone, we also performed MLM on the other three tasks. Although the model with experiment as the only fixed factor is the best model, the analysis of variance for the alertness * experiment model still shows a reliable effect of alertness on threshold (*F*_(1,68)_ = 5.032, *p* = 0.028). Therefore, we observe a common effect of alertness level on threshold of detection and discrimination decisions though it is not necessarily visible in each individual experiment.

**Table 3:** Model comparison for threshold.

| Model | Parameter | AIC | BIC | Log-likelihood | $p > (x^2)$ |
| --- | --- | --- | --- | --- | --- |
| Null | Fixed: mean, Random: participant ID | 481.19 | 490.45 | -237.59 | - |
| Alertness | Fixed: alertness,<br>Random: participant ID | 473.76 | 486.11 | -232.88 | 0.002 ** |
| Experiment | Fixed: experiment,<br>Random: participant ID | 425.69 | 444.21 | -206.84 | 2.81e-13 *** |
| Alertness * Experiment | Fixed: alertness * experiment,<br>Random: participant ID | 420.07 | 450.94 | -200.03 | 1.35e-13 *** |

\*\* $p < 0.01$ ; \*\*\* $p < 0.001$ .

**Table 4:** Type III analysis of variance table for the alertness * experiment model on threshold.

| Model elements | Sum Sq | Mean Sq | NumDF | DenDF | $F$ | $P(> F)$ |
| --- | --- | --- | --- | --- | --- | --- |
| Alertness | 4.628 | 4.628 | 1 | 81 | 12.859 | 5.73e-4 *** |
| Experiment | 33.139 | 11.046 | 3 | 81 | 30.692 | 2.36e-13 *** |
| Alertness:experiment | 1.549 | 0.516 | 3 | 81 | 1.435 | 0.239 |

\*\*\* $p < 0.001$ .

Like in the case of slope, we also performed the analysis using the three other methods to split alertness. The results are mostly consistent with the micro-measures method: alertness * experiment is the winning model (Table S10) for three methods except for omissions in which the alertness * experiment model improves marginally compared to the experiment model (*Chi – Squared* = 8.976, *p* = 0.062). Further ANOVA analysis shows that both main effects of alertness and experiment are reliable across different methods(Table S11). The only difference is that there is a reliable interaction effect under the RTs split (*F*_(3,89)_ = 8.866, *p* < 0.001), which comes from strong effects for the auditory masking task and TMS task as suggested by the *post hoc* analysis (Table S12). In addition, we checked the influence of parameters’ specifications for the curve fitting, and found that the main effects of alertness and experiment are reliable for both group-best parameters and strict parameters, consistent with our main findings (Supplementary Section 6).

### 3.3 Alertness affects perceptual sensitivity, but not criterion

To further characterize the modulation exerted by fluctuations of alertness on perceptual decision-making, we applied SDT to complete the testing of hypotheses H2 and H3 defined in the preregistration. We assessed the effects of alertness on perceptual sensitivity (d’) and the criterion (C) extracted from SDT analysis for each participant, and found that d’ decreases while C remains unchanged as the alertness level goes lower.

**Table 5:**
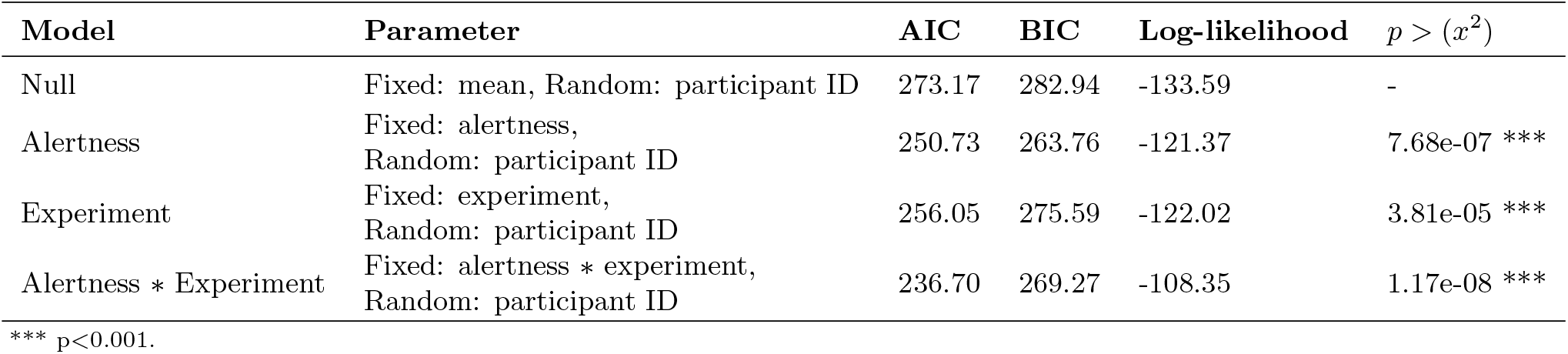
Model comparison for d’.

**Table 6:** Type III analysis of variance table for the alertness * experiment model on d’.

| Model elements | Sum Sq | Mean Sq | NumDF | DenDF | $F$ | $P(> F)$ |
| --- | --- | --- | --- | --- | --- | --- |
| Alertness | 4.189 | 4.189 | 1 | 96 | 25.171 | 2.41e-06 *** |
| Experiment | 4.416 | 1.472 | 3 | 96 | 8.846 | 3.1e-05 *** |
| Alertness:experiment | 0.439 | 0.146 | 3 | 96 | 0.88 | 0.455 |

\*\*\* $p < 0.001$ .

In detail, the comparison of the distribution of d’ is shown in Fig. 5. Compared to the high alertness level, the distributions of d’ tend to be lower and more variable in low alertness. There is a small effect in detection tasks and a medium to large effect in discrimination tasks. The variance of d’ increases in three tasks (*F*_(19,19)_ = 0.293, *p* = 0.01 for Exp. 1, *F*_(26,26)_ = 0.234, *p* < 0.001 for Exp. 3, and *F*_(32,32)_ = 0.3, *p* = 0.001 for Exp. 4) except for the TMS task (*F*_(15,15)_ = 0.76, *p* = 0.602). In MLM, the alertness * experiment is the winning model (Table 5), and its ANOVA results show that both alertness (*F*_(1,96)_ = 25.171, *p* < 0.001) and experiment (*F*_(3,96)_ = 8.846, *p* < 0.001) have a strong effect on d’ (Table 6), indicating that there is a decrease in sensitivity as people get drowsy regardless of experiment types. There was no evidence for an interaction effect with d’ (*F*_(3,96)_ = 0.88, *p* = 0.455), which is different from what we found with slope of the sigmoidal curve. This implies that alertness modulates detection and discrimination differentially when looking at how precise we are in taking a decision but not for an overall measure of performance like perceptual sensitivity(d’).

**Figure 5:**
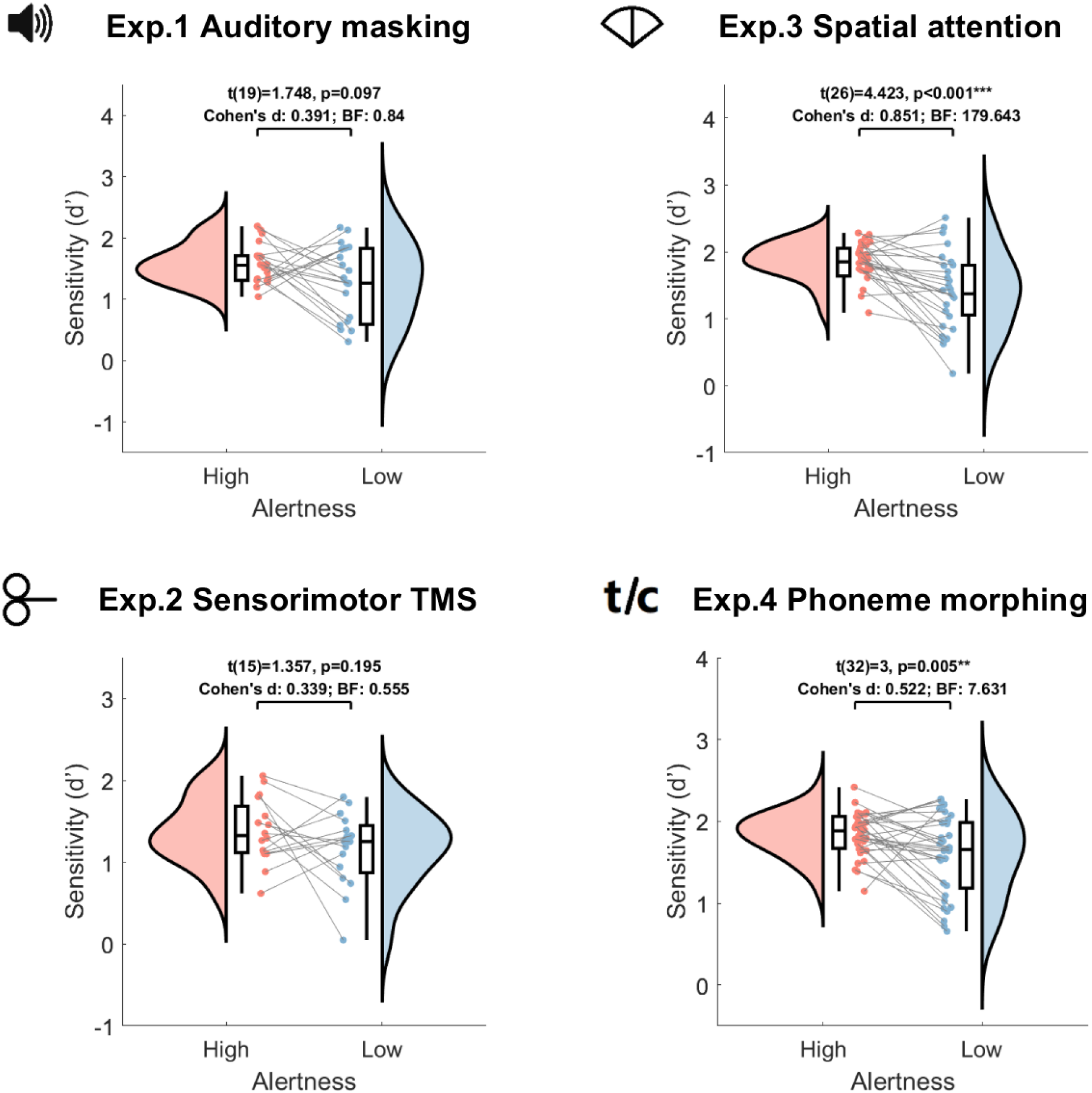
Distributions of d prime. Distributions of d’ are shown for each task and each alertness level (defined by the micro-measures method). Results of the t-test, effect size and the Bayes factor are shown in the figure. d’ decreases reliably with alertness level for spatial attention and phoneme discrimination tasks. Although there are no obvious effects in detection tasks, the MLM including four experiments found a reliable effect of alertness and no interaction effect of alertness and experimental settings (see text and Table 5 - 6).

As with slope and threshold, the analyses of d’ using the complementary categorizing methods confirmed the conver-gence of the results obtained with the micro-measures split. Alertness * experiment is the winning model for all different splitting methods (Table S16). Further ANOVA analysis of the winning model (Table S17) shows consistent outcomes to micro-measures splitting that alertness has a strong effect on d’ while it shows no interaction with experiment for RTs and omissions splitting methods. For the *θ/α* ratio method, there is a weak interaction effect between alertness and experiment on d’ (*F*_(3,107)_ = 2.707, *p* = 0.049). The experiment factor also shows a reliable main effect on d’ for all three complementary splitting methods. To sum up, these complementary analyses confirm that d’ decreases with lower alertness for all four alertness level categorization methods and it shows no interaction with experiment settings for most categorization methods.

The other parameter of SDT, criterion, shows little change from wakefulness to low alertness (Fig. 6). Results from MLM analysis show that the experiment model has the smallest values of AIC and BIC and the second largest loglikelihood value (Table 7). Comparison between the experiment model and the alertness * experiment model shows that including alertness in the model does not significantly improve the fit (*Chi – Squared* = 2.404, *p* = 0.662). ANOVA outcome of the alertness * experiment model (Table 8) suggests no effect for alertness (*F*_(1,94)_ = 0.558, *p* = 0.457) and no interaction effect between alertness and experiment (*F*_(3,94)_ = 0.438, *p* = 0.727), while the main effect of experiment is reliable (*F*_(3,94)_ = 10.323, *p* < 0.001). These results indicate that the criterion to make a decision is unlikely to be influenced by alertness level.

**Table 7:** Model comparison for C.

| Model | Parameter | AIC | BIC | Log-likelihood | $p > (x^2)$ |
| --- | --- | --- | --- | --- | --- |
| Null | Fixed: mean, Random: participant ID | 105.5 | 115.21 | -49.749 | - |
| Alertness | Fixed: alertness,<br>Random: participant ID | 106.4 | 119.34 | -49.199 | 0.294 |
| Experiment | Fixed: experiment,<br>Random: participant ID | 84.73 | 104.15 | -36.365 | 6.58e-06 *** |
| Alertness * Experiment | Fixed: alertness * experiment,<br>Random: participant ID | 90.325 | 122.69 | -35.163 | 1.35e-4 *** |

\*\*\* $p < 0.001$ .

**Table 8:** Type III analysis of variance table for the alertness * experiment model on C.

| Model elements | Sum Sq | Mean Sq | NumDF | DenDF | $F$ | $P(> F)$ |
| --- | --- | --- | --- | --- | --- | --- |
| Alertness | 0.038 | 0.038 | 1 | 94 | 0.558 | 0.457 |
| Experiment | 2.098 | 0.699 | 3 | 94 | 10.323 | 6.16e-06 *** |
| Alertness:experiment | 0.089 | 0.03 | 3 | 94 | 0.438 | 0.727 |

\*\*\* $p < 0.001$ .

**Figure 6:**
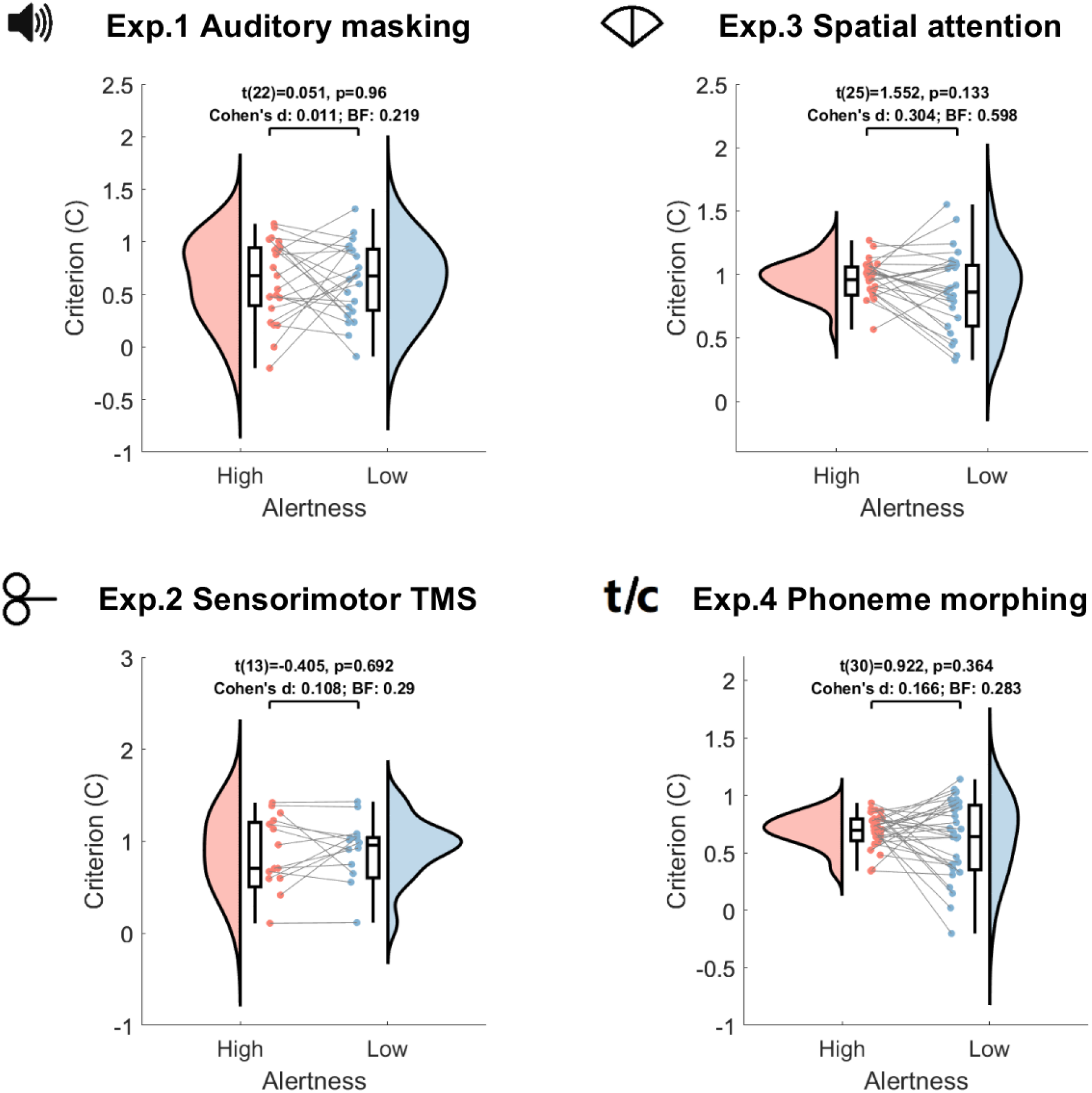
Distributions of criterion. Distributions of criterion are shown for each task and each alertness level (defined by the micro-measures method). Results of the t-test, effect size and the Bayes factor are shown in the figure. Criterion shows no reliable change for all tasks.

The same analysis on criterion using *θ/α* splitting has consistent results as the micro-measures method (Table S19 - S21). However, alertness shows a strong effect on criterion for the RTs and omissions methods (Table S20). In addition to that, there is a reliable interaction effect (*F*_(3,103_) = 2.699, *p* = 0.05) between alertness and experiment when using the RTs as the splitting method. *Post hoc* analysis (Table S21) reveals that criterion decreases as people enter a low alertness state for the spatial attention task when low alertness is defined by RTs and omissions, and for the phoneme morphing discrimination task when low alertness is defined by omissions. Therefore, the alertness level categorized by pre-stimulus brain state has little effect on criterion, while lower alertness defined by post-event behavioural markers significantly influences the criterion.

To summarize the results, the decision-making performance follows a sigmoidal curve in both high and low alertness states, but with different parameter characteristics of slope and threshold. The main findings show lower detection and discrimination sensitivity to stimuli in low alertness with a shallower slope of the psychometric curve and lower perceptual sensitivity (d’). This decrease in sensitivity is more pronounced for discrimination than for detection decisions as captured by the degree of change in the slope of the curve. Threshold shows a small increase with lower alertness and there is no change in the criterion to make the decision during fluctuations of alertness defined by pre-stimulus brain state.

## 4 Discussion

Despite the pervasiveness of the alertness fluctuations happening in every human being throughout the day, the underlying mechanisms of how it affects perceptual decision-making and cognitive functions are surprisingly understudied. Our study demonstrates that there is an overwhelming agreement on how alertness fluctuations modulate perceptual decision-making. Here we developed an analysis framework using SDT and psychophysics to look at relative differences between high alertness and metastable state of low alertness in two detection and two discrimination decision-making tasks. We showed that the slope of the sigmoidal curve decreases with alertness, associated with lower accuracy of sensory representation (Gold and Ding, 2013). Complementary, we observed a subtle increase in threshold, indicating that participants may require more sensory evidence with lower alertness. Although the increase is small and only evident in the kinaesthetic detection experiment (TMS). In terms of general performance, the d’, capturing perceptual sensitivity, reveals a systematic and convergent decrease for all tasks, while the criterion, a reflection of internal rules to take a decision, may not vary. The changes observed in these four parameters, capturing complementary aspects of cognition, characterize the effects of alertness fluctuations on perceptual detection and discrimination in humans.

### 4.1 Decreased decisions’ precision with lower alertness may reflect a re-balance of the internal noise of the neural systems

First, we found that performance in low alertness follows a similar sigmoidal psychometric curve as in the high alertness state but the curve is shallower (smaller slope) with decreasing alertness. Slope of the psychometric curve describes changes in response accuracy as opposed to varying stimulus levels (Gold and Ding, 2013) and is often considered a reflection of the level of sensory noise (Morgan et al., 2012; Whiteley and Sahani, 2008). A decrease in slope with lower alertness was also found in previous research using the same datasets of auditory detection, TMS-induced kinaesthetic detection and spatial discrimination that were analysed separately (Jagannathan et al., 2018; Noreika, Canales-Johnson, et al., 2020; Noreika et al., 2017). Here we found that the phenomenon is common across different experiment settings using a general analytical framework, indicating a diminished ability to precisely distinguish similar stimuli for both detection and discrimination tasks as people become less alert. This decreased precision of decision may reflect the increased variance in the underlying neural processes during the transition from wakefulness to lower alertness (Marreiros et al., 2008; Noreika, Canales-Johnson, et al., 2020). In other words, people become less sensitive to exogenous noise (weak stimulus strength or introduction of another stimulus) as their endogenous noise increases (lower alertness level) in detection and discrimination decision-making. In addition, by combining four experiments together, here we surprisingly found a distinction showing the decrease in slope of the psychometric curve accompanied by lower alertness is larger for discrimination than for detection decisions. It indicates that fluctuations in the inner noise have a stronger effect on the precision of discrimination decisions than detection decision-making.

The change in the precision of decision is also found in association with attention, which has been conceptualized as a measure of the inner noise of the system when integration of information is needed (Chennu et al., 2016; Lu and Dosher, 1998). Attention reduces the neural noise that is taken into account for the decision-making process, increasing the signal-to-noise ratio, and sensitivity as a result (Nunez et al., 2017; Smith and Ratcliff, 2009). Cohen and Maunsell (M. R. Cohen and Maunsell, 2011) studied how uncontrolled attentional fluctuations affect behavioural performance and found that the ability to detect minor changes in stimuli is considerably enhanced by attention. Herrmann and colleagues (Herrmann et al., 2010) also found that attention could increase the slope of the psychometric function. Alertness and attention have an intertwined relationship. One key component of attention, sustained attention, is not only integrated with cognitive processing but also connected to the alertness systems (Oken et al., 2006; Posner, 2008). The fluctuations of the neural dynamics underlying the perceptual and cognitive systems created by the transition of consciousness from high alertness to low alertness might show convergent effects to those proposed for sustained attention (Bareham et al., 2014), contributing in a similar manner to the changes in decision-making processes.

The interaction between noise and decision complexity has been proposed previously (Smith and Ratcliff, 2009). When external noise is introduced to the stimulus, its effect on performance is small in detection, but large in discrimination tasks (Smith and Wolfgang, 2007; Smith et al., 2004). Convergently, here we found an interaction between the inner noise (alertness level) and task types showing that the precision of decision declines more with low alertness for discrimination than detection. As far as the nature of the tasks is concerned, discrimination (distinguishing between two stimuli) has more decision complexity than detection (detecting one stimulus) (Sagi and Julesz, 1984; Smith and Ratcliff, 2009), and more complex neural networks may be involved in the former than in the latter (Correa et al., 2004). This makes discrimination easier to carry out as reflected by the steeper slope of the psychometric curve in discrimination than detection tasks. When alertness level declines, the increased inner noise may impair more information processing involved in discrimination as more neural networks may be affected, hence the decrease in the precision of decision (slope) is greater for discrimination than detection.

### 4.2 Increased threshold with lower alertness may suggest compensation for less efficient information processing

Next, the position of the psychometric curve is captured by threshold, which we found increased as alertness declines not systematically for each task but reliably in the aggregated data. When looking into the experiments individually, we found only a reliable change in threshold for the TMS sensorimotor task and not enough evidence of a shifted threshold for the other three tasks. This result confirms previous findings from the same datasets of auditory mask and TMS sensorimotor task (Noreika, Canales-Johnson, et al., 2020; Noreika et al., 2017). However, when combining all experiments together, we could see a clear effect of alertness level on threshold regardless of the experiment type. The effect is not necessarily visible in every single experiment but needs aggregated data. Since threshold represents the strength of evidence where one alternative exceeds the other (Macmillan and Creelman, 2004), it appears that participants require more evidence of the target in lower alertness to reach the same performance level as compared to full wakefulness. Threshold is also a reflection of the decision rule towards one alternative over another (Gold and Ding, 2013). Our result of the increased threshold indicates a propensity towards stronger target stimuli induced by a lower alertness level in decision-making tasks. Such a shift in decision rule has been considered as resulting from cognitive resource constraints such as time and memory capacity (Bossaerts and Murawski, 2017). Here the decreased alertness level results in cognitive and perceptual resource constraints and further prevents people from achieving optimal performance.

### 4.3 Perceptual representations have broader and closer distributions with lower alertness

Further analysis of SDT allows us to disentangle perceptual sensitivity and decision-making criteria during the transition from wakefulness to lower alertness. Our results show that perceptual sensitivity (d’) decreases as alertness level declines, which is consistent with previous research (Jagannathan et al., 2022). d’ is considered to reflect the underlying decision variable which can be represented as a Gaussian distribution for each alternative in decision-making (Gold and Ding, 2013). A smaller d’ could indicate an increase in their common standard deviation and a decrease in the difference between the means of the Gaussian distributions. The former, increased standard deviation, represents that the inner noise gets larger which makes it more difficult to distinguish between the two. The latter represents that the perceptual representations of two alternatives are closer to each other which could also impair perceptual sensitivity. Either way, the decreased perception at sleep onset could be attributed to the diminished transmission of external information to the cortex, or its integration (Goupil and Bekinschtein, 2012). In addition, contrary to the change of slope, the variation of d’ is higher with low alertness in three tasks. It suggests that the perceptual sensitivity may become more dispersed among the population as people enter a lower alertness state, while the precision of decision (sensitivity to distinguish similar stimuli at the behavioural level) tends to be more convergent, for the group, in lower alertness compared to wakefulness. Furthermore, different from sensitivity to the stimuli (slope), the decrease in perceptual sensitivity with lower alertness shows no discrepancy between detection and discrimination tasks. It indicates that alertness level may comparably affect the sensitivity of detection and discrimination at the general performance level across different task conditions (measured by a normalized metric d’), but differentially at the local level around the threshold of the curve for difficult trials that are most indistinguishable (measured by slope).

On the other hand, we found no evidence for a difference in criterion of SDT in alertness regardless of the decision type. This result extends previous findings in the spatial attention task to wider decision-making scenarios (Jagannathan et al., 2022). Criterion in decision-making is a rule that the observer uses to partition the underlying distributions of stimuli (Macmillan and Creelman, 2004). The lack of evidence for a change in criterion in varied alertness may indicate a stable subjective rule while the sensory processing ability diminishes with lower alertness. Based on the relation between SDT and the psychometric curve, we can see that the criterion to make decisions and the distance between representation distributions of the noise and the stimulus (or distributions of two stimuli in discrimination) contribute jointly to the threshold of performance. Since there is no evidence for a change in criterion at different alertness levels, the small increase in threshold we found should not be attributed to the change in criterion. Instead, it could be caused by a decrease in the distance between distributions of alternatives with lower alertness. Therefore, there are indeed changes in response propensity (captured by threshold) due to changes in distribution distances which criterion (C) does not reflect. The smaller distance in lower alertness is also consistent with our finding of a decreased perceptual sensitivity (d’, computed as the difference in distance of distributions divided by the common standard deviation), although a small decrease in the distance could only account for part of the decline in sensitivity; the change in sensitivity also comes from the increased noise (standard deviation of the distribution) that is reported here.

### 4.4 The effects of alertness on decision making are convergent between different alertness measures and fitting parameters

We did several variations of the analysis to validate our results. For the alertness level categorization, we applied both EEG and behavioural measures to categorize alertness levels, to capture the different aspects of low alertness. Micro-measures method and *θ/α* ratio are both pretrial measures defined before the sensory perturbation and based on EEG/neural signals, reflecting a physiological ready state for decision-making (Samaha et al., 2020). On the other hand, split methods of RTs and proportion of omissions are measures of behaviour as the outcome of the perceptual decision (Goupil and Bekinschtein, 2012; Ogilvie et al., 1989). There is a consistency between pretrial measures and post-event behavioural measures, and both seem to point towards clear metastability of the arousal state *per se* and convergent on its modulation to cognitive performance except for criterion in SDT, as shown by the MLM results on aggregated data.

Alertness levels are fairly stable in neighbouring trials suggesting a clear hysteresis of the transition of consciousness even when perturbated systematically by tones, noise, words or tactile stimuli. Furthermore, previously our neural methods to separate different alertness states can only be applied when the participants have their eyes closed (Jagannathan et al., 2022). In current results, it seems that the type of perceptual decision does not influence the alertness metrics that we calculated, nor neural or behavioural, which is good news to extend the alertness-modulating cognition framework beyond auditory and tactile experiments. The convergence of effects of alertness on decision-making parameters seen between neural and behavioural methods to define high and low alertness periods is encouraging, and may help extend the separation between these states of the transition to other cases that involve visual or cross-modal integration since the behavioural methods are comparable even if the neural methods cannot be applied.

### 4.5 Implications of the results

The influence of alertness fluctuations on cognitive functions is insufficiently investigated despite its pervasiveness and wide application. This maybe is due to the fact the fields of cognitive neuroscience and health and industrial-organizational (I/O) psychology have been historically separated (Butler and Senior, 2007). While cognitive neuroscience deals with the characterization of the underlying mechanisms of thought and action, applied psychology relates to human performance like driving behaviour and shift work. Applied psychology has been concentrating on capturing errors that can lead to accidents with little regard for neurocognitive mechanisms (Schreier et al., 2018). It is key to bring those willing to explain and those willing to predict real-life events together to create a truly translational programme of research that can use the neurocognitive models to predict performance in a wide set of contexts and events. Here we offer an example of the connection between performance and underlying cognitive factors in a common context of varying alertness levels, shedding light on both the explanation and prediction of perceptual decision-making.

Our convergent framework consists of four methods to categorize alertness levels capturing both pre-trial neural characteristics and post-event behavioural indicators and variations of psychometric curve-fitting specifications. The framework provides robust and reliable results of changes in decision-making performance in the transition from wakefulness to low alertness. In addition, the combination of four detection and discrimination experiments allows for testing the effect of alertness level across different modalities and experimental settings and offers a comparison between detection and discrimination decision-making. Furthermore, the joint application of psychometric function fitting and SDT connects behavioural performance to the underlying decision process, revealing that it is the broader and closer distributions of perceptual representation that lead to impaired performance of decision-making while the relative criterion remains stable from high alertness to low alertness. In this paper, this integrated analysis framework proves effective and promising for future use to illustrate how alertness level affects perceptual decision-making.

Moving forward, this research opens up new avenues for future investigation. To start, the relationship among different methods of alertness classification could be further explored. Here we found convergence regarding performance outcomes, and it would be informative to see the degree of agreement these methods show. Considerable overlap of alertness categorization is expected, and the features of those most consistent trials could be further explored with the potential to develop an integrated and comprehensive classification method for alertness levels. Further, current evidence of increased internal noise and demand for stronger stimuli at lower alertness levels suggests changes in information processing during the transition between alertness levels. Future works would benefit from applying information theory tools to explore the underlying information dynamics of decision-making.

## Footnotes

1 In a subset of the discrimination data, the target and noise stimuli were shifted and tested, showing no difference from current results for all four behavioural measures in the phoneme morphing task and for three measures in the spatial attention task except for criterion. The criterion in the spatial attention task decreased with lower alertness when using right-sided stimuli as the target and it was most likely due to the left bias.

## Notes

### Competing Interest Statement

The authors have declared no competing interest.

