## Supplementary material for "Effects of alertness on perceptual detection and discrimination"

### 1 Psychometric curves with different splits

Figure S1 **Curve fitting using different splits**

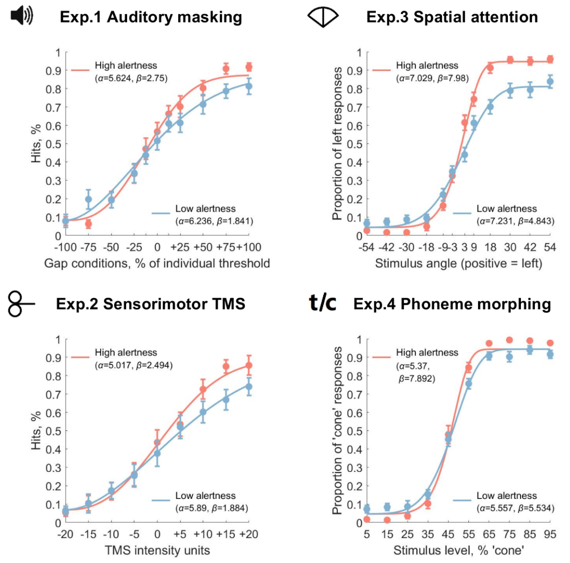


(a) **Split by micro-measures**

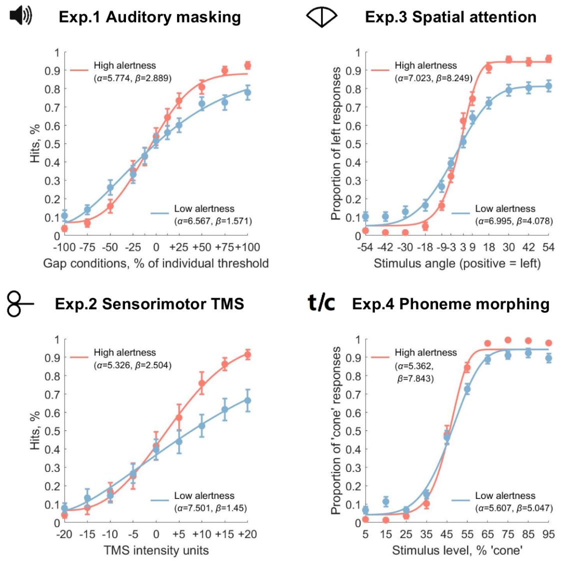


(c) **Split by RTs**

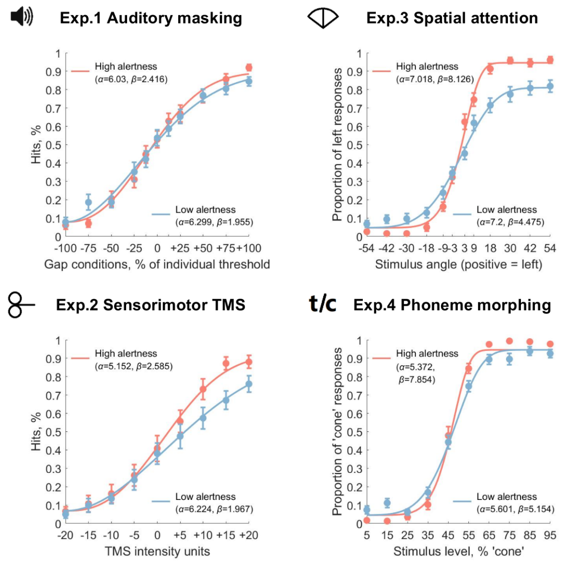


(b) **Split by $\theta/\alpha$ ratio**

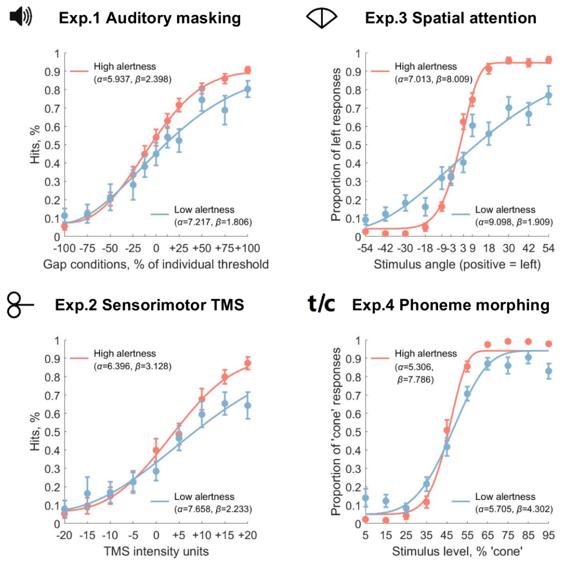


(d) **Split by Omissions**

Fitted Weibull curves for four methods of alertness level classification. (a) Split by micro-measures is shown in the main text. (b-d) Split by other classification methods shows consistent patterns as micro-measures: slope decreases and threshold increases in lower alertness compared to the high alertness state (see the following MLM results).

#### 2 Slope with different splits

Figure S2 **Distributions of slope using different splits**

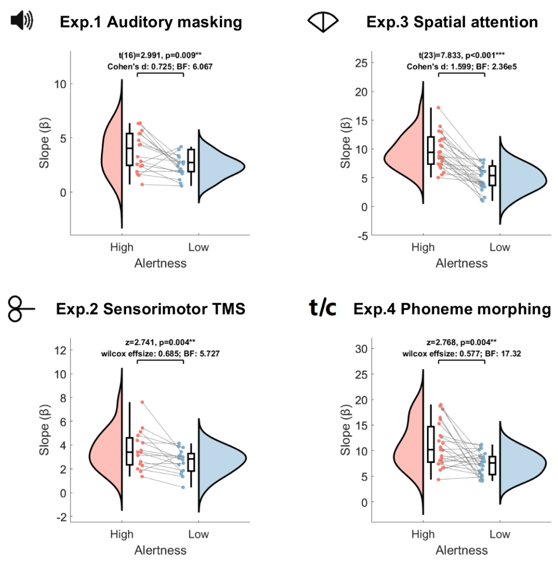


(a) **Split by micro-measures**

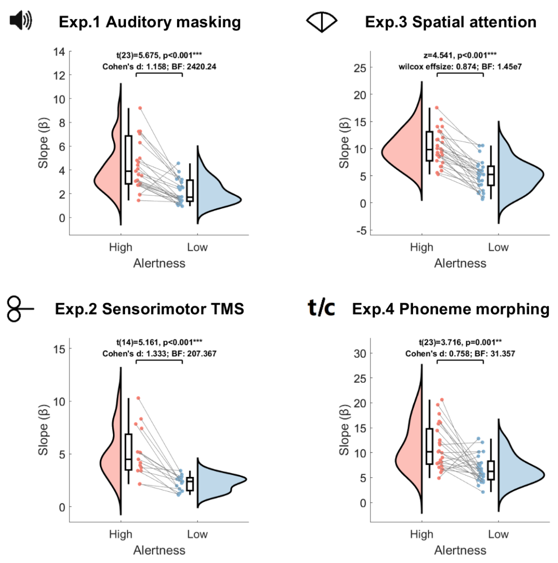


(c) **Split by RTs**

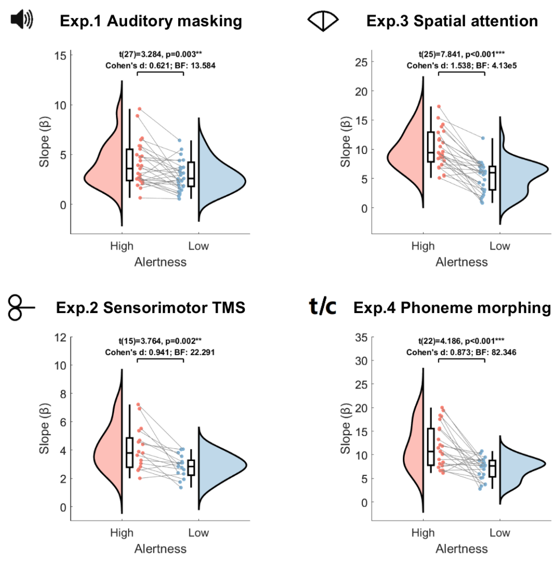


(b) **Split by $\theta/\alpha$ ratio**

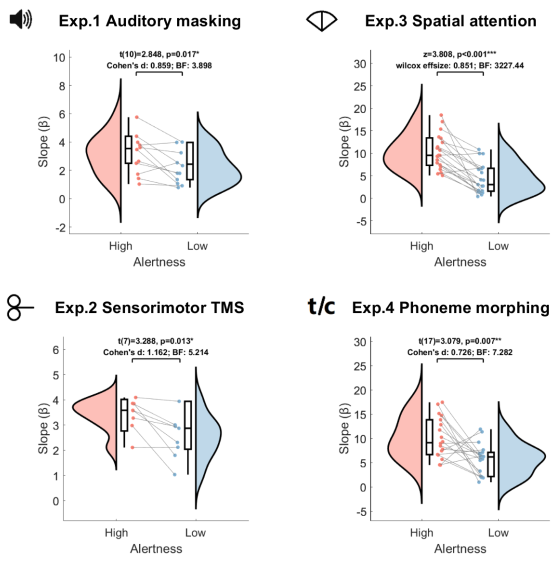


(d) **Split by Omissions**

Distributions of slope for four methods of alertness level classification. (a) Split by micro-measures is shown in the main text. (b-d) Split by other classification methods shows a consistent decrease in slope in lower alertness compared to the high alertness state.

Table S1 **Model comparison for slope**

| Model | Parameter | Micro-measures | | $\theta/\alpha$ ratio | | RTs | | Omissions | |
|---|---|---|---|---|---|---|---|---|---|
| | | Log-likelihood | $p > (x^2)$ | Log-likelihood | $p > (x^2)$ | Log-likelihood | $p > (x^2)$ | Log-likelihood | $p > (x^2)$ |
| Null | Fixed: mean, Random: participant ID | -444.20 | - | -519.31 | - | -509.71 | - | -328.87 | - |
| Alertness | Fixed: alertness, Random: participant ID | -424.03 | 2.14e-10 *** | -495.19 | 3.77e-12 *** | -479.02 | 4.73e-15 *** | -314.58 | 8.99e-08 *** |
| Experiment | Fixed: experiment, Random: participant ID | -400.37 | < 2.2e-16 *** | -474.25 | < 2.2e-16 *** | -471.97 | 2.87e-16 *** | -309.94 | 3.03e-08 *** |
| Alertness ∗ Experiment | Fixed: alertness ∗ experiment, Random: participant ID | -368.61 | < 2.2e-16 *** | -437.29 | < 2.2e-16 *** | -431.63 | < 2.2e-16 *** | -285.77 | 7.42e-16 *** |

Table S2 **Type III analysis of variance table for the alertness ∗ experiment model on slope**

| Model elements | Micro-measures | | $\theta/\alpha$ ratio | | RTs | | Omissions | |
|---|---|---|---|---|---|---|---|---|
| | $F$ value | $Pr(> F)$ | $F$ value | $Pr(> F)$ | $F$ value | $Pr(> F)$ | $F$ value | $Pr(> F)$ |
| Alertness | 51.228 | 3.53e-10 *** | 72.582 | 2.76e-13 *** | 88.305 | 5.13e-15 *** | 26.338 | 3.6e-06 *** |
| Experiment | 60.006 | < 2.2e-16 *** | 53.584 | < 2.2e-16 *** | 46.147 | < 2.2e-16 *** | 22.808 | 7.92e-10 *** |
| Alertness:experiment | 6.12 | 8.42e-4 *** | 8.464 | 4.97e-05 *** | 3.886 | 0.012 * | 4.731 | 0.005 ** |

Table S3 ***Post hoc* comparisons for slope**

| Low alertness - high alertness | Micro-measures | | $\theta/\alpha$ ratio | | RTs | | Omissions | |
|---|---|---|---|---|---|---|---|---|
| | $z$ value | $Pr(\|z\|)$ | $z$ value | $Pr(\|z\|)$ | $z$ value | $Pr(\|z\|)$ | $z$ value | $Pr(\|z\|)$ |
| Auditory masking detection | -1.62 | 0.737 | -1.963 | 0.502 | -2.755 | 0.105 | -0.76 | 0.995 |
| TMS sensorimotor detection | -1.172 | 0.94 | -1.573 | 0.763 | -3.132 | 0.037 * | -0.616 | 0.999 |
| Auditory spatial discrimination | -7.026 | < 0.001 *** | -7.222 | < 0.001 *** | -7.472 | < 0.001 *** | -6.275 | < 0.001 *** |
| Auditory phoneme discrimination | -5.43 | < 0.001 *** | -6.885 | < 0.001 *** | -6.149 | < 0.001 *** | -4.496 | < 0.001 *** |

### 3 Slope with drowsy session only

One factor potentially influencing data selection for high and low alertness states comes from differences in the structure of the experimental design. Since the discrimination tasks have an extra awake session while in the detection tasks, high and low alertness are extracted from the same session, the stronger effect for discrimination may come from the differences in the nature of high alertness trials created by the independent session. To test this possibility, we applied the same analysis only on drowsy sessions for discrimination tasks. This means that instead of using the awake session, we use relatively high alertness trials in the drowsy session as high alertness data while keeping relatively low alertness trials in the drowsy session as low alertness data. As a result, we find that alertness ∗ experiment is still the winning model (Table S4), and the effect of alertness remains strong while the interaction effect disappears for three out of four types of splits (Table S5). *Post hoc* analysis tells us that the alertness effect is still larger in the discrimination tasks than the detection tasks generally (Table S6).

Table S4 **Model comparison for slope with drowsy session only**

| Model | Parameter | Micro-measures | | $\theta/\alpha$ ratio | | RTs | | Omissions | |
|---|---|---|---|---|---|---|---|---|---|
| | | Log-likelihood | $p > (x^2)$ | Log-likelihood | $p > (x^2)$ | Log-likelihood | $p > (x^2)$ | Log-likelihood | $p > (x^2)$ |
| Null | Fixed: mean, Random: participant ID | -343.99 | - | -460.43 | - | -500.56 | - | -328.81 | - |
| Alertness | Fixed: alertness, Random: participant ID | -330.24 | 1.56e-07 *** | -441.51 | 7.67e-10 *** | -471.55 | 2.61e-14 *** | -317.29 | 1.6e-06 *** |
| Experiment | Fixed: experiment, Random: participant ID | -312.44 | 1.28e-13 *** | -424.84 | 2.39e-15 *** | -470.64 | 6.38e-13 *** | -312.44 | 3.66e-07 *** |
| Alertness ∗ Experiment | Fixed: alertness*experiment, Random: participant ID | -295.86 | < 2.2e-16 *** | -401.34 | < 2.2e-16 *** | -438.06 | < 2.2e-16 *** | -298.37 | 1.01e-10 *** |

Table S5 **Type III analysis of variance table for the alertness ∗ experiment model on slope with drowsy session only**

| Model elements | Micro-measures | | $\theta/\alpha$ ratio | | RTs | | Omissions | |
|---|---|---|---|---|---|---|---|---|
| | $F$ value | $Pr(> F)$ | $F$ value | $Pr(> F)$ | $F$ value | $Pr(> F)$ | $F$ value | $Pr(> F)$ |
| Alertness | 35.663 | 9.52e-08 *** | 44.794 | 1.75e-09 *** | 80.955 | 3.48e-14 *** | 18.588 | 6.25e-05 *** |
| Experiment | 34.665 | 1.04e-13 *** | 35.982 | 2e-15 *** | 28.567 | 4.55e-13 *** | 14.587 | 3.22e-07 *** |
| Alertness:experiment | 1.962 | 0.128 | 3.213 | 0.027 * | 2.346 | 0.078 . | 1.778 | 0.161 |

Table S6 ***Post hoc* comparisons for slope with drowsy session only**

| Low alertness - high alertness | Micro-measures | | $\theta/\alpha$ ratio | | RTs | | Omissions | |
|---|---|---|---|---|---|---|---|---|
| | $z$ value | $Pr(\|z\|)$ | $z$ value | $Pr(\|z\|)$ | $z$ value | $Pr(\|z\|)$ | $z$ value | $Pr(\|z\|)$ |
| Auditory masking detection | -2.054 | 0.44 | -2.494 | 0.19 | -2.938 | 0.063 . | -0.969 | 0.975 |
| TMS sensorimotor detection | -1.486 | 0.811 | -1.998 | 0.47 | -3.341 | 0.018 * | -0.786 | 0.993 |
| Auditory spatial discrimination | -4.495 | < 0.001 *** | -6.498 | < 0.001 *** | -5.687 | < 0.001 *** | -3.937 | 0.002 ** |
| Auditory phoneme discrimination | -4.013 | 0.001 ** | -3.016 | 0.05 * | -6.537 | < 0.001 *** | -4.357 | < 0.001 *** |

### 4 Slope with other fittings

Another possible confounding factor is the parameters used in psychometric curve fitting. For the main analysis, we use individual best guess rate ($\gamma$) and lapse rate ($\lambda$) when fitting the sigmoidal curve to get a better fitting for each participant and to capture the group variability. In order to exclude the influence of these two parameters on the difference of slope between states and test whether the results still hold when considering fewer individual variances, we fit the curve using a pair of group best $\gamma$ and $\lambda$ rates for participants in one experiment. The results for the group parameters (Table S7 - S9) demonstrate that the individual variances in the curve fitting do not impair our main conclusion that alertness level has a common effect on slope across different experiments. In addition, to compare different detection and discrimination experiments, we also apply a more strict criterion to fit the curve, which is using 0.05 as both guess rate and lapse rate for all participants across four experiments. The results (Table S7 - S9) again indicate that parameter selection of the curve fitting has little influence on our results.

Table S7 **Model comparison for slope with other fittings**

| Model | Parameter | Micro-measures | | $\theta/\alpha$ ratio | | RTs | | Omissions | |
|---|---|---|---|---|---|---|---|---|---|
| | | Log-likelihood | $p > (x^2)$ | Log-likelihood | $p > (x^2)$ | Log-likelihood | $p > (x^2)$ | Log-likelihood | $p > (x^2)$ |
| **Group fitting** | | | | | | | | | |
| Null | Fixed: mean, Random: participant ID | -462.7 | - | -531.07 | - | -532.18 | - | -376.5 | - |
| Alertness | Fixed: alertness, Random: participant ID | -443.09 | 3.81e-10 *** | -512.16 | 7.82e-10 *** | -503.58 | 3.91e-14 *** | -360.85 | 2.2e-08 *** |
| Experiment | Fixed: experiment, Random: participant ID | -433.94 | 2e-12 *** | -500.48 | 3.32e-13 *** | -503.95 | 3.34e-12 *** | -356.82 | 1.45e-08 *** |
| Alertness * Experiment | Fixed: alertness * experiment, Random: participant ID | -405.34 | < 2.2e-16 *** | -468.51 | < 2.2e-16 *** | -467.55 | < 2.2e-16 *** | -325.36 | < 2.2e-16 *** |
| **Strict fitting** | | | | | | | | | |
| Null | Fixed: mean, Random: participant ID | -475.31 | - | -528.53 | - | -530.73 | - | -366.42 | - |
| Alertness | Fixed: alertness, Random: participant ID | -451.78 | 6.86e-12 *** | -506.77 | 4.22e-11 *** | -501.26 | 1.62e-14 *** | -349.33 | 5.03e-09 *** |
| Experiment | Fixed: experiment, Random: participant ID | -441.19 | 1.02e-14 *** | -490.03 | < 2.2e-16 *** | -499.00 | 1.07e-13 *** | -346.59 | 1.26e-08 *** |
| Alertness * Experiment | Fixed: alertness * experiment, Random: participant ID | -400.58 | < 2.2e-16 *** | -448.51 | < 2.2e-16 *** | -452.24 | < 2.2e-16 *** | -312.88 | < 2.2e-16 *** |

Table S8 **Type III analysis of variance table for the alertness * experiment model on slope with other fittings**

| Model elements | Micro-measures | | $\theta/\alpha$ ratio | | RTs | | Omissions | |
|---|---|---|---|---|---|---|---|---|
| | $F$ value | $Pr(> F)$ | $F$ value | $Pr(> F)$ | $F$ value | $Pr(> F)$ | $F$ value | $Pr(> F)$ |
| **Group fitting** | | | | | | | | |
| Alertness | 47.333 | 1.15e-09 *** | 54.718 | 6.33e-11 *** | 70.577 | 6.5e-13 *** | 37.193 | 7.21e-08 *** |
| Experiment | 27.921 | 1.7e-12 *** | 28.953 | 2.85e-13 *** | 26.651 | 2.16e-12 *** | 24.462 | 1.29e-10 *** |
| Alertness:experiment | 6.715 | 4.2e-4 *** | 10.083 | 8.29e-06 *** | 5.451 | 0.002 ** | 8.925 | 5.17e-05 *** |
| **Strict fitting** | | | | | | | | |
| Alertness | 65.018 | 5.23e-12 *** | 69.759 | 7.11e-13 *** | 85.32 | 1.106e-14 *** | 39.37 | 4.02e-08 *** |
| Experiment | 36.33 | 4.9e-15 *** | 42.363 | < 2.2e-16 *** | 35.747 | 2.63e-15 *** | 25.497 | 8.19e-11 *** |
| Alertness:experiment | 13.573 | 2.85e-07 *** | 15.215 | 4.2e-08 *** | 10.684 | 4.48e-06 *** | 9.475 | 3.15e-05 *** |

Table S9 ***Post hoc* comparisons for slope with other fittings**

| **Low alertness - high alertness** | **Micro-measures** | | **$\theta/\alpha$ ratio** | | **RTs** | | **Omissions** | |
|---|---|---|---|---|---|---|---|---|
| | $z$ **value** | $Pr(\|z\|)$ | $z$ **value** | $Pr(\|z\|)$ | $z$ **value** | $Pr(\|z\|)$ | $z$ **value** | $Pr(\|z\|)$ |
| **Group fitting** | | | | | | | | |
| Auditory masking detection | -2.299 | 0.283 | -0.996 | 0.973 | -2.743 | 0.105 | -0.199 | 1 |
| TMS sensorimotor detection | -0.274 | 1 | -1.145 | 0.942 | -1.895 | 0.541 | -0.943 | 0.982 |
| Auditory spatial discrimination | -7.137 | < 0.001 *** | -7.724 | < 0.001 *** | -8.439 | < 0.001 *** | -7.681 | < 0.001 *** |
| Auditory phoneme discrimination | -5.179 | < 0.001 *** | -5.515 | < 0.001 *** | -5.388 | < 0.001 *** | -4.964 | < 0.001 *** |
| **Strict fitting** | | | | | | | | |
| Auditory masking detection | -1.713 | 0.666 | -1.158 | 0.941 | -2.559 | 0.166 | -0.473 | 1 |
| TMS sensorimotor detection | -0.585 | 0.999 | -1.022 | 0.97 | -2.207 | 0.339 | -0.663 | 0.998 |
| Auditory spatial discrimination | -10.106 | < 0.001 *** | -9.731 | < 0.001 *** | -10.366 | < 0.001 *** | -8.102 | < 0.001 *** |
| Auditory phoneme discrimination | -5.52 | < 0.001 *** | -5.686 | < 0.001 *** | -5.108 | < 0.001 *** | -5.413 | < 0.001 *** |

### 5 Threshold with different splits

Figure S3 **Distributions of threshold using different splits**

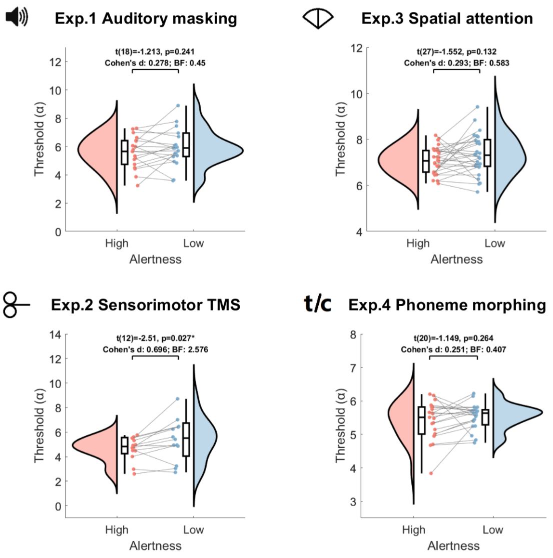


(a) **Split by micro-measures**

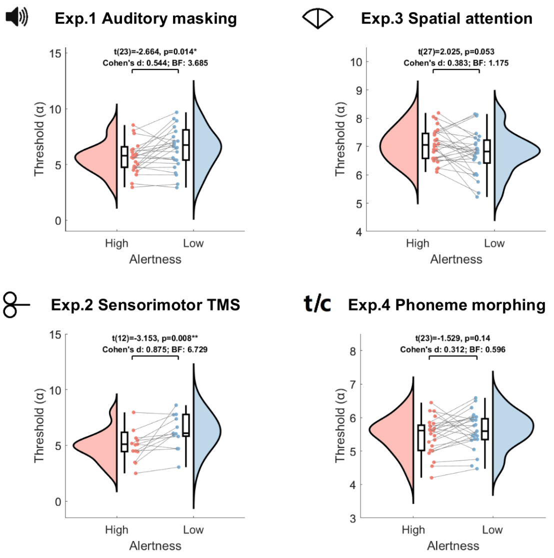


(c) **Split by RTs**

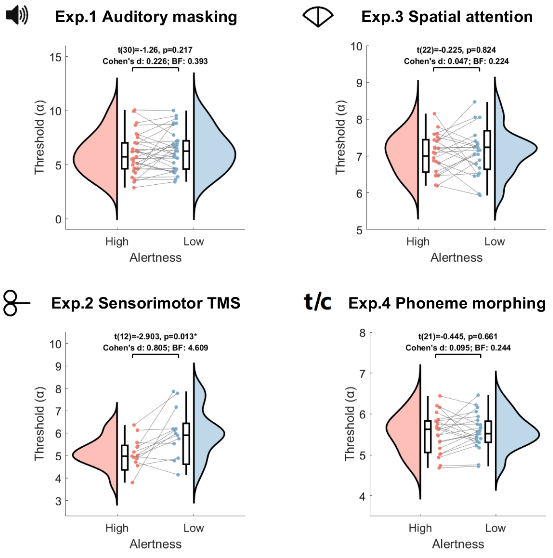


(b) **Split by $\theta/\alpha$ ratio**

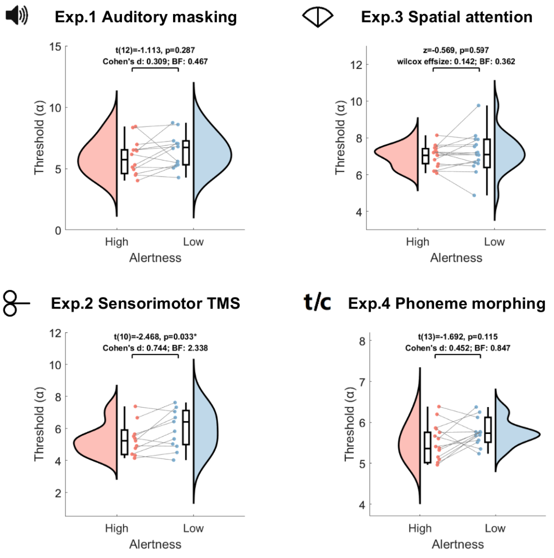


(d) **Split by Omissions**

Distributions of threshold for four methods of alertness level classification. (a) Split by micro-measures is shown in the main text. (b-d) Split by other classification methods shows an evident increase in threshold in lower alertness compared to the high alertness state in the TMS experiment but not for other experiments.

Table S10 **Model comparison for threshold**

| Model | Parameter | Micro-measures | | $\theta/\alpha$ ratio | | RTs | | Omissions | |
|---|---|---|---|---|---|---|---|---|---|
| | | Log-likelihood | $p > (x^2)$ | Log-likelihood | $p > (x^2)$ | Log-likelihood | $p > (x^2)$ | Log-likelihood | $p > (x^2)$ |
| Null | Fixed: mean, Random: participant ID | -237.59 | - | -279.67 | - | -279.33 | - | -151.12 | - |
| Alertness | Fixed: alertness, Random: participant ID | -232.88 | 0.002 ** | -277.04 | 0.022 * | -275.45 | 0.005 ** | -147.57 | 0.008 ** |
| Experiment | Fixed: experiment, Random: participant ID | -206.84 | 2.81e-13 *** | -267.47 | 2.05e-05 *** | -264.56 | 1.73e-06 *** | -139.30 | 2.96e-05 *** |
| Alertness ∗ Experiment | Fixed: alertness ∗ experiment, Random: participant ID | -200.03 | 1.35e-13 *** | -261.45 | 5.99e-06 *** | -249.05 | 1.17e-10 *** | -134.81 | 3.12e-05 *** |

Table S11 **Type III analysis of variance table for the alertness ∗ experiment model on threshold**

| Model elements | Micro-measures | | $\theta/\alpha$ ratio | | RTs | | Omissions | |
|---|---|---|---|---|---|---|---|---|
| | $F$ value | $Pr(> F)$ | $F$ value | $Pr(> F)$ | $F$ value | $Pr(> F)$ | $F$ value | $Pr(> F)$ |
| Alertness | 12.859 | 5.73e-4 *** | 7.886 | 0.006 ** | 18.408 | 4.52e-05 *** | 8.73 | 0.005 ** |
| Experiment | 30.692 | 2.36e-13 *** | 9.363 | 1.92e-05 *** | 11.673 | 1.6e-06 *** | 9.889 | 2.68e-05 *** |
| Alertness:experiment | 1.435 | 0.239 | 2.341 | 0.079 . | 8.866 | 3.33e-05 *** | 0.639 | 0.593 |

Table S12 ***Post hoc* comparisons for threshold**

| Low alertness - high alertness | Micro-measures | | $\theta/\alpha$ ratio | | RTs | | Omissions | |
|---|---|---|---|---|---|---|---|---|
| | $z$ value | $Pr(\|z\|)$ | $z$ value | $Pr(\|z\|)$ | $z$ value | $Pr(\|z\|)$ | $z$ value | $Pr(\|z\|)$ |
| Auditory masking detection | 1.53 | 0.768 | 1.837 | 0.558 | 3.714 | 0.005 ** | 1.724 | 0.641 |
| TMS sensorimotor detection | 3.103 | 0.036 * | 3.074 | 0.038 * | 4.548 | < 0.001 *** | 2.292 | 0.272 |
| Auditory spatial discrimination | 1.545 | 0.759 | 0.138 | 1 | -1.447 | 0.817 | 0.826 | 0.99 |
| Auditory phoneme discrimination | 0.692 | 0.997 | 0.181 | 1 | 0.752 | 0.994 | 0.926 | 0.98 |

### 6 Threshold with other fittings

When using different settings for guess rate ($\gamma$) and lapse rate ($\lambda$), alertness ∗ experiment model on threshold is still the winning model (Table S13) for most categorization methods except for omissions in group fitting in which alertness ∗ experiment model improves marginally compared to the experiment model ($Chi - Squared = 9.328, p = 0.053$). The main effects of alertness and experiment are reliable (Table S14) for both group best parameters and the strict parameters, which is consistent with previous findings. We find a significant interaction effect under micro-measures, $\theta/\alpha$ ratio and RTs split when using group parameters and strict parameters. *Post hoc* analysis (Table S15) shows that the interaction effect is not necessarily driven by differences between detection and discrimination tasks. We find that alertness has a strong influence on threshold in the TMS experiment across the first three methods as the results for individual best parameters.

Table S13 **Model comparison for threshold with other fittings**

| Model | Parameter | Micro-measures | | $\theta/\alpha$ ratio | | RTs | | Omissions | |
|---|---|---|---|---|---|---|---|---|---|
| | | Log-likelihood | $p > (x^2)$ | Log-likelihood | $p > (x^2)$ | Log-likelihood | $p > (x^2)$ | Log-likelihood | $p > (x^2)$ |
| **Group fitting** | | | | | | | | | |
| Null | Fixed: mean, Random: participant ID | -238.61 | - | -237.89 | - | -308.93 | - | -226.79 | - |
| Alertness | Fixed: alertness, Random: participant ID | -234.47 | 0.004 ** | -235.69 | 0.036 * | -302.97 | 5.54e-4 *** | -224.26 | 0.025 * |
| Experiment | Fixed: experiment, Random: participant ID | -229.27 | 3.19e-4 *** | -226.04 | 2.9e-05 *** | -301.77 | 0.002 ** | -218.04 | 5.59e-4 *** |
| Alertness ∗ Experiment | Fixed: alertness ∗ experiment, Random: participant ID | -220.74 | 8.1e-06 *** | -218.66 | 2.49e-06 *** | -288.41 | 7.96e-07 *** | -213.38 | 3.59e-4 *** |
| **Strict fitting** | | | | | | | | | |
| Null | Fixed: mean, Random: participant ID | -249.05 | - | -297.77 | - | -354.41 | - | -231.48 | - |
| Alertness | Fixed: alertness, Random: participant ID | -237.42 | 1.42e-06 *** | -289.46 | 4.55e-05 *** | -342.13 | 7.2e-07 *** | -228.47 | 0.014 * |
| Experiment | Fixed: experiment, Random: participant ID | -236.21 | 1.11e-05 *** | -285.43 | 1.79e-05 *** | -347.38 | 0.003 ** | -222.98 | 7.07e-4 *** |
| Alertness ∗ Experiment | Fixed: alertness ∗ experiment, Random: participant ID | -219.79 | 3e-10 *** | -272.75 | 1.41e-08 *** | -329.92 | 2.3e-08 *** | -217.43 | 2.11e-4 *** |

Table S14 **Type III analysis of variance table for the alertness $*$ experiment model on threshold with other fittings**

| Model elements | Micro-measures | | $\theta/\alpha$ ratio | | RTs | | Omissions | |
|---|---|---|---|---|---|---|---|---|
| | $F$ value | $Pr(>F)$ | $F$ value | $Pr(>F)$ | $F$ value | $Pr(>F)$ | $F$ value | $Pr(>F)$ |
| **Group fitting** | | | | | | | | |
| Alertness | 10.203 | 0.002 ** | 7.701 | 0.007 ** | 21.025 | 1.59e-05 *** | 5.308 | 0.025 * |
| Experiment | 7.069 | 3.01e-4 *** | 9.285 | 2.7e-05 *** | 5.212 | 0.002 ** | 6.771 | 5.23e-4 *** |
| Alertness:experiment | 3.113 | 0.031 * | 3.707 | 0.015 * | 5.396 | 0.002 ** | 1.476 | 0.23 |
| **Strict fitting** | | | | | | | | |
| Alertness | 30.121 | 5.43e-07 *** | 21.404 | 1.34e-05 *** | 32.487 | 1.61e-07 *** | 6.187 | 0.016 * |
| Experiment | 10.232 | 1.03e-05 *** | 9.567 | 1.68e-05 *** | 5.081 | 0.003 ** | 6.551 | 6.62e-4 *** |
| Alertness:experiment | 3.409 | 0.022 * | 3.068 | 0.032 * | 3.668 | 0.015 * | 1.77 | 0.163 |

Table S15 ***Post hoc*** **comparisons for threshold with other fittings**

| Low alertness - high alertness | Micro-measures | | $\theta/\alpha$ ratio | | RTs | | Omissions | |
|---|---|---|---|---|---|---|---|---|
| | $z$ value | $Pr(\|z\|)$ | $z$ value | $Pr(\|z\|)$ | $z$ value | $Pr(\|z\|)$ | $z$ value | $Pr(\|z\|)$ |
| **Group fitting** | | | | | | | | |
| Auditory masking detection | 3.042 | 0.039 * | 0.899 | 0.985 | 2.139 | 0.372 | 0.006 | 1 |
| TMS sensorimotor detection | 3.144 | 0.028 * | 3.921 | 0.002 ** | 5.1 | $< 0.001$ *** | 2.358 | 0.257 |
| Auditory spatial discrimination | 0.162 | 1 | 0.347 | 1 | 0.386 | 1 | 2.119 | 0.396 |
| Auditory phoneme discrimination | -0.01 | 1 | -0.109 | 1 | 0.88 | 0.986 | 0.205 | 1 |
| **Strict fitting** | | | | | | | | |
| Auditory masking detection | 3.554 | 0.008 ** | 2.282 | 0.278 | 4.276 | $< 0.001$ *** | -0.055 | 1 |
| TMS sensorimotor detection | 3.722 | 0.004 ** | 3.763 | 0.004 ** | 4.335 | $< 0.001$ *** | 2.39 | 0.241 |
| Auditory spatial discrimination | 3.852 | 0.003 ** | 3.199 | 0.027 * | 2.389 | 0.232 | 2.533 | 0.177 |
| Auditory phoneme discrimination | 0.02 | 1 | -0.031 | 1 | 0.436 | 1 | 0.245 | 1 |

#### 7 d’ with different splits

Figure S4 **Distributions of d’ using different splits**

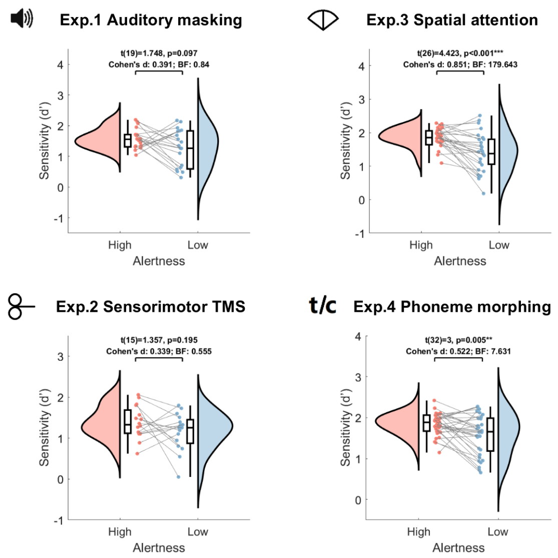


(a) **Split by micro-measures**

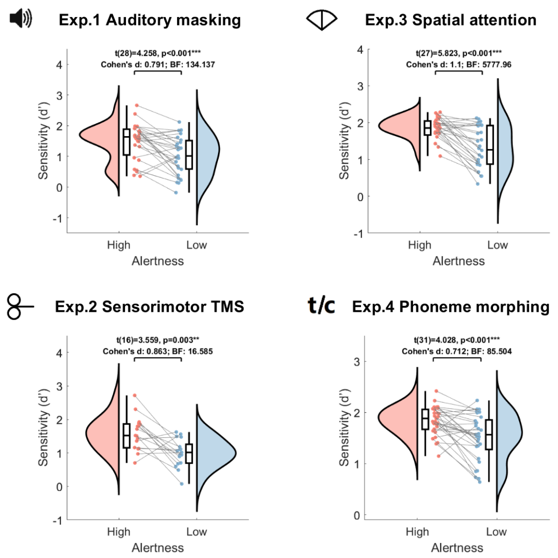


(c) **Split by RTs**

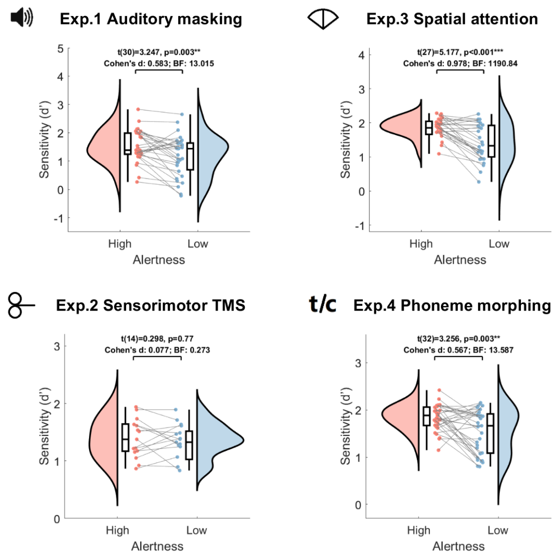


(b) **Split by $\theta/\alpha$ ratio**

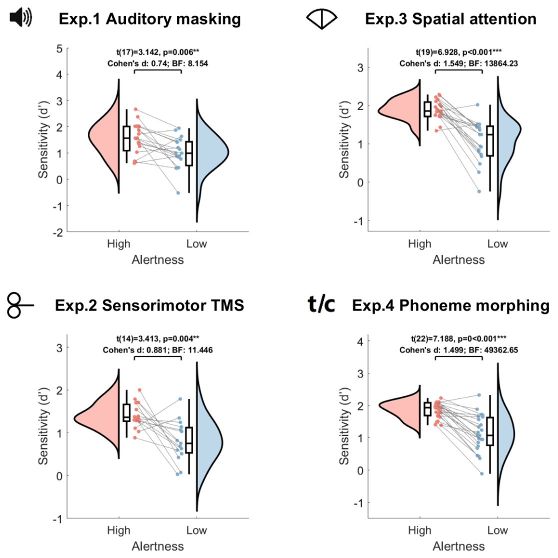


(d) **Split by Omissions**

Distributions of d prime for four methods of alertness level classification. (a) Split by micro-measures is shown in the main text. (b-d) Split by other classification methods shows a consistent decrease in d prime in lower alertness compared to the high alertness state except for the TMS task split by $\theta/\alpha$ ratio.

Table S16 **Model comparison for d'**

| **Model** | **Parameter** | **Micro-measures** | | **$\theta/\alpha$ ratio** | | **RTs** | | **Omissions** | |
|---|---|---|---|---|---|---|---|---|---|
| | | **Log-likelihood** | $p > (x^2)$ | **Log-likelihood** | $p > (x^2)$ | **Log-likelihood** | $p > (x^2)$ | **Log-likelihood** | $p > (x^2)$ |
| Null | Fixed: mean,<br>Random: participant ID | -133.59 | - | -156.85 | - | -175 | - | -136.16 | - |
| Alertness | Fixed: alertness,<br>Random: participant ID | -121.37 | 7.68e-07 *** | -139.75 | 4.98e-09 *** | -146.01 | 2.63e-14 *** | -102.96 | 3.68e-16 *** |
| Experiment | Fixed: experiment,<br>Random: participant ID | -122.02 | 3.81e-05 *** | -149.27 | 0.002 ** | -163.45 | 3.85e-05 *** | -130.95 | 0.015 * |
| Alertness $*$ Experiment | Fixed: alertness $*$ experiment,<br>Random: participant ID | -108.35 | 1.17e-08 *** | -128.26 | 5.49e-10 *** | -132.29 | 1.07e-15 *** | -94.952 | 4.41e-15 *** |

Table S17 **Type III analysis of variance table for the alertness $*$ experiment model on d'**

| **Model elements** | **Micro-measures** | | **$\theta/\alpha$ ratio** | | **RTs** | | **Omissions** | |
|---|---|---|---|---|---|---|---|---|
| | $F$ **value** | $Pr(> F)$ | $F$ **value** | $Pr(> F)$ | $F$ **value** | $Pr(> F)$ | $F$ **value** | $Pr(> F)$ |
| Alertness | 25.171 | 2.41e-06 *** | 31.349 | 1.68e-07 *** | 81.449 | 8.76e-15 *** | 96.9 | 3.28e-15 *** |
| Experiment | 8.846 | 3.1e-05 *** | 5.43 | 0.002 ** | 8.604 | 3.65e-05 *** | 4.981 | 0.003 ** |
| Alertness:experiment | 0.88 | 0.455 | 2.707 | 0.049 * | 1.474 | 0.226 | 0.805 | 0.495 |

Table S18 ***Post hoc* comparisons for d'**

| **Low alertness - high alertness** | **Micro-measures** | | **$\theta/\alpha$ ratio** | | **RTs** | | **Omissions** | |
|---|---|---|---|---|---|---|---|---|
| | $z$ **value** | $Pr(\|z\|)$ | $z$ **value** | $Pr(\|z\|)$ | $z$ **value** | $Pr(\|z\|)$ | $z$ **value** | $Pr(\|z\|)$ |
| Auditory masking detection | -2.213 | 0.339 | -3.855 | 0.003 ** | -4.726 | < 0.001 *** | -4.006 | 0.002 ** |
| TMS sensorimotor detection | -1.542 | 0.781 | -0.175 | 1 | -4.519 | < 0.001 *** | -3.887 | 0.003 ** |
| Auditory spatial discrimination | -4.185 | < 0.01 *** | -5.013 | < 0.001 *** | -5.579 | < 0.001 *** | -6.236 | < 0.001 *** |
| Auditory phoneme discrimination | -2.539 | 0.177 | -3.384 | 0.015 * | -3.309 | 0.02 * | -5.897 | < 0.001 *** |

### 8 Criterion with different splits

Figure S5 **Distributions of Criterion using different splits**

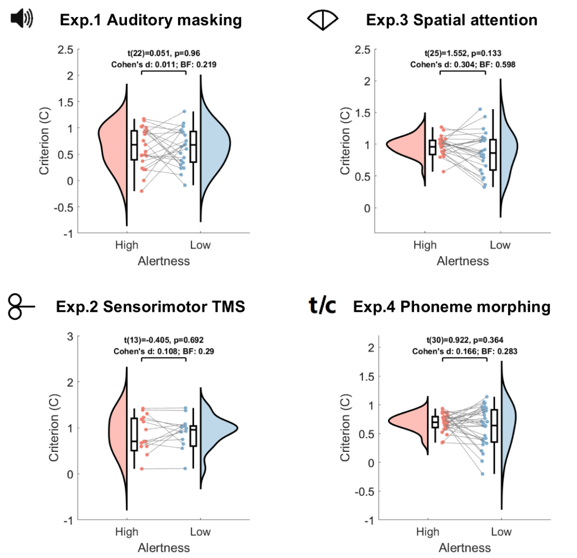


(a) **Split by micro-measures**

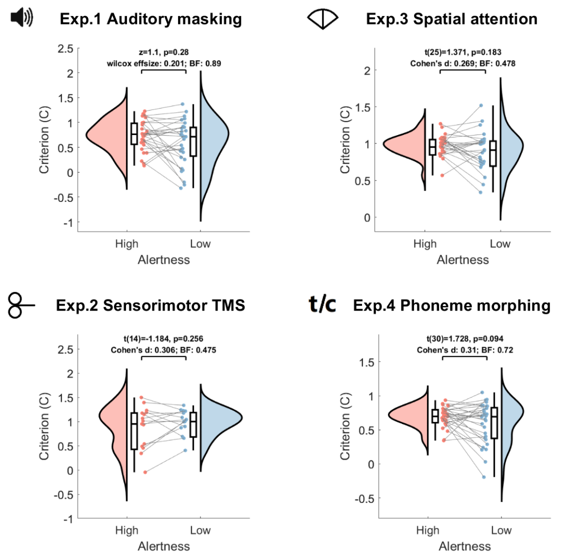


(b) **Split by $\theta/\alpha$ ratio**

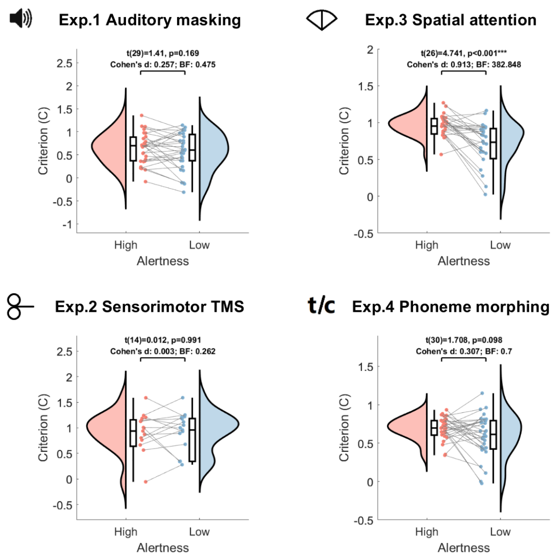


(c) **Split by RTs**

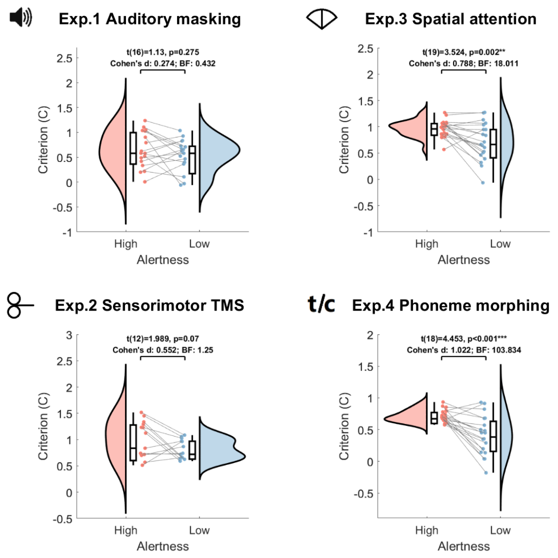


(d) **Split by Omissions**

Distributions of criterion for four methods of alertness level classification. (a) Split by micro-measures is shown in the main text. (b-d) Split by other classification methods show no change in criterion in different alertness levels except for the spatial attention task split by RTs and omissions, and the phoneme morphing discrimination task split by omissions.

Table S19 **Model comparison for criterion**

| Model | Parameter | Micro-measures | | $\theta/\alpha$ ratio | | RTs | | Omissions | |
|---|---|---|---|---|---|---|---|---|---|
| | | Log-likelihood | $p > (x^2)$ | Log-likelihood | $p > (x^2)$ | Log-likelihood | $p > (x^2)$ | Log-likelihood | $p > (x^2)$ |
| Null | Fixed: mean,<br>Random: participant ID | -49.749 | - | -52.069 | - | -46.884 | - | -41.197 | - |
| Alertness | Fixed: alertness,<br>Random: participant ID | -49.199 | 0.294 | -49.692 | 0.029 * | -40.064 | 2.21e-4 *** | -28.931 | 7.31e-07 *** |
| Experiment | Fixed: experiment,<br>Random: participant ID | -36.365 | 6.58e-06 *** | -37.002 | 1.29e-06 *** | -34.531 | 1.78e-05 *** | -28.246 | 1e-05 *** |
| Alertness ∗ Experiment | Fixed: alertness ∗ experiment,<br>Random: participant ID | -35.163 | 1.35e-4 *** | -31.872 | 1.06e-06 *** | -23.814 | 8.21e-08 *** | -14.253 | 2.48e-09 *** |

Table S20 **Type III analysis of variance table for the alertness ∗ experiment model on criterion**

| Model elements | Micro-measures | | $\theta/\alpha$ ratio | | RTs | | Omissions | |
|---|---|---|---|---|---|---|---|---|
| | $F$ value | $Pr(> F)$ | $F$ value | $Pr(> F)$ | $F$ value | $Pr(> F)$ | $F$ value | $Pr(> F)$ |
| Alertness | 0.558 | 0.457 | 2.342 | 0.129 | 11.846 | 8.36e-4 *** | 28.856 | 9.94e-07 *** |
| Experiment | 10.323 | 6.16e-06 *** | 11.686 | 1.21e-06 *** | 9.307 | 1.68e-05 *** | 10.478 | 9.17e-06 *** |
| Alertness:experiment | 0.438 | 0.727 | 1.886 | 0.137 | 2.699 | 0.05 * | 1.181 | 0.324 |

Table S21 ***Post hoc* comparisons for criterion**

| Low alertness - high alertness | Micro-measures | | $\theta/\alpha$ ratio | | RTs | | Omissions | |
|---|---|---|---|---|---|---|---|---|
| | $z$ value | $Pr(\|z\|)$ | $z$ value | $Pr(\|z\|)$ | $z$ value | $Pr(\|z\|)$ | $z$ value | $Pr(\|z\|)$ |
| Auditory masking detection | -0.069 | 1 | -2.443 | 0.212 | -1.482 | 0.803 | -1.305 | 0.894 |
| TMS sensorimotor detection | 0.353 | 1 | 1.153 | 0.941 | -0.013 | 1 | -2.242 | 0.32 |
| Auditory spatial discrimination | -1.216 | 0.925 | -1.051 | 0.964 | -4.315 | < 0.001 *** | -3.401 | 0.015 * |
| Auditory phoneme discrimination | -0.909 | 0.985 | -1.547 | 0.772 | -1.736 | 0.643 | -4.027 | 0.001 ** |
